# Bacteriophage protein Dap2 inhibits bacterial type III secretion system and synergizes with Dap1 to evade anti-phage immunity

**DOI:** 10.1101/2025.03.13.642734

**Authors:** Jingru Zhao, Yuhao Zhu, Chenchen Wang, Fan Tian, Jun Deng, Jianglin Liao, Zhuojun Zhong, Jiazhen Liu, Nannan Guo, Shuai Le, Haihua Liang

## Abstract

The evolutionary arms race between bacterial immunity and phages has driven the emergence of sophisticated anti-defense systems (ADSs). However, certain ADSs exhibit incomplete suppression of their cognate defense systems, suggesting functional cooperation between multiple ADSs targeting the same bacterial safeguard. In this study, we characterize Dap2, a protein encoded by a *Pseudomonas aeruginosa* phage PaoP5, which directly binds to the Lon protease to prevent the degradation of the phage-encoded HNH endonuclease. Deletion of *dap2* in PaoP5 exhibits significantly impaired genome packaging due to insufficient levels of HNH. Strikingly, Dap2 synergizes with its genomically adjacent partner Dap1, a previously identified HNH-binding protein providing partial Lon resistance, to achieve complete protection of HNH. Beyond anti-defense activity, Dap2 disrupts host virulence by sequestering the type III secretion system (T3SS) transcriptional activator ExsA, suppressing bacterial pathogenicity while redirecting metabolic resources toward phage progeny production. This study unveils a dual functional ADS that simultaneously modulates bacterial virulence and anti-phage immunity, both aimed at ensuring phage survival and maximizing progeny production. Furthermore, it elucidates a novel mechanism whereby phages employ synergistic ADS pairs (Dap1/Dap2) to achieve complete neutralization of a bacterial defense system, when individual components provide only partial protection. These findings significantly enhance our understanding of the intricate evolutionary arms race between phages and their bacterial hosts.

## Introduction

The coevolutionary arms race between bacteriophages and their bacterial hosts over billions of years has driven the emergence of diverse bacterial immune mechanisms, including CRISPR-Cas, restriction-modification (RM) systems, CBASS, Pycsar, Gabija, and Thoeris^1–4^. In response, phages have evolved sophisticated anti-defense systems (ADSs) to counteract these immune strategies. While numerous ADSs targeting CRISPR-Cas^5,6^ and RM systems^7,8^ have been well characterized, recent studies have uncovered novel phage-encoded countermeasures against CBASS, Pycsar, Gabija, and Thoeris pathways^9–12^.

Characterized ADSs employ a variety of molecular strategies to neutralize host immunity, such as direct inhibition of defense proteins through binding, enzymatic modification or deactivation of immune components, degradation or sequestration of signaling molecules, metabolic compensation for host-induced depletion of essential molecules, and structural camouflage of phage components to evade detection^10,13^. To date, approximately 150 distinct ADSs have been functionally characterized^10^. However, the vast diversity of bacterial defense mechanisms necessitates ongoing discovery and mechanistic analysis of ADSs^14,15^. Notably, some ADSs confer only partial resistance to bacterial defenses, as illustrated by the phage protein Acb2: its induction only partially rescues phage titers in JBD67*Δacb2* and JBD18 variants^16^. This suggests that phages may employ multiple counterstrategies against individual immune systems to enhance protection^10^.

Phage reproduction relies heavily on host resources, prompting phages to frequently manipulate bacterial transcription, DNA replication, translation, and cell division pathways to optimize infection cycles^17–19^. These host-modulating factors, typically expressed during early infection phases, represent potential targets for novel antimicrobial therapies^17^. While current research has predominantly focused on phage-encoded lethal proteins^18,20^, the regulatory elements affecting bacterial virulence and their direct biological benefits for phages remain poorly understood^21,22^.

Here we establish a systematic framework for identifying virulence-modulating hypothetical proteins in *P. aeruginosa* phage PaoP5, with a particular focus on regulators of the type 3 secretion system (T3SS), a critical virulence determinant^23,24^. We identified the phage protein Dap2, which directly binds to ExsA^25^, the master regulator of T3SS, suppressing T3SS expression and bacterial pathogenicity. This suppression likely redirects host resources toward phage progeny production. Intriguingly, Dap2 also interacts with the bacterial Lon protease to prevent degradation of the phage-encoded HNH endonuclease^26^. Deletion of *dap2* results in impaired genome packaging and reduced progeny viability. Our findings complement the previously characterized Dap1 protein, which stabilizes HNH through direct binding^22^. The coordinated action of Dap1 and Dap2 establishes a dual protection mechanism against Lon-mediated antiviral defense: Dap1 physically shields HNH, while Dap2 inhibits protease activity. This synergistic system ensures complete protection of the essential HNH endonuclease from host degradation.

This study highlights the dual functionality of phage ADS in simultaneously regulating bacterial virulence and countering host immunity, both aimed at maximizing phage progeny production. Furthermore, it reveals that phages can deploy synergistic anti-defense proteins (Dap1/Dap2) with distinct mechanisms to neutralize single bacterial defense systems, expanding our understanding of phage evolutionary strategies in the ongoing arms race with their hosts.

## Results

### Identification of a phage protein Dap2 that inhibits T3SS

To systematically identify phage-encoded inhibitors of *Pseudomonas aeruginosa* type III secretion system (T3SS), a critical virulence apparatus for host cell invasion^23,27^, we developed a dual-plasmid fluorescence reporter assay (Fig. 1a). The transcriptional activity of T3SS was monitored by fusing the *exsC* promoter (encoding a core T3SS structural protein) to *luxCDABE* biosensor genes in plasmid pMS402^28^. Concurrently, 51 PaoP5 open reading frames (ORFs) that do not affect bacterial growth by expressing them individually from IPTG-inducible expression vector pME6032 (Table S1-S2). Upon co-transformation of both plasmids into PAO1, fluorescence attenuation served as a proxy for T3SS inhibition. Interestingly, IPTG-induced expression of ORF004 (herein designated *Dap2*, Defense anti-phage protein 2) demonstrated dose-dependent suppression of luminescence (Fig. S1a-b), suggesting specific targeting of T3SS regulatory components. To further validate the inhibitory effect of Dap2 on T3SS functionality, we analyzed the promoter activity of key T3SS effector genes (*exoS*, *exoY*, and *exoT*) in WT PAO1 versus PAO1 harboring pME6032-*dap2*. The results demonstrated that Dap2 overexpression significantly suppressed promoter activity across all tested genes (Fig. S1c). Consistent with these transcriptional changes, immunoblot analysis revealed a marked reduction in ExoS protein levels and ExsA (the master T3SS regulator) expression in the *dap2*-overexpressing strain compared to the vector control (Fig. 1b). These findings collectively establish Dap2 as a potent T3SS suppressor and warrant detailed investigation into its molecular mechanism of action. To gain insight into how Dap1 affects T3SS, we performed RNA sequencing (RNA-seq) comparing the transcriptional profiles of PAO1/p-*dap2* (*dap2*-overexpressing strain) and PAO1 carrying the empty vector during exponential growth. Overexpression of *dap2* altered the expression of 229 genes (|fold change| > 2, *P* < 0.05), with 93 upregulated and 136 downregulated genes (Fig. 1c). Strikingly, KEGG pathway enrichment analysis identified the type III secretion system as the most significantly affected pathway, with multiple T3SS-associated genes clustering in this category (Fig. S2a-b; Table S3). To confirm the RNA-seq data reliability, we selected four randomly chosen genes (*hemH*, *fhp*, *pcrv*, *pscF*) and five T3SS-related genes for qRT-PCR validation. The qRT-PCR results exhibited strong concordance with RNA-seq trends, showing consistent upregulation or downregulation patterns in response to *dap2* overexpression (Fig. S2c).

**Fig. 1.**
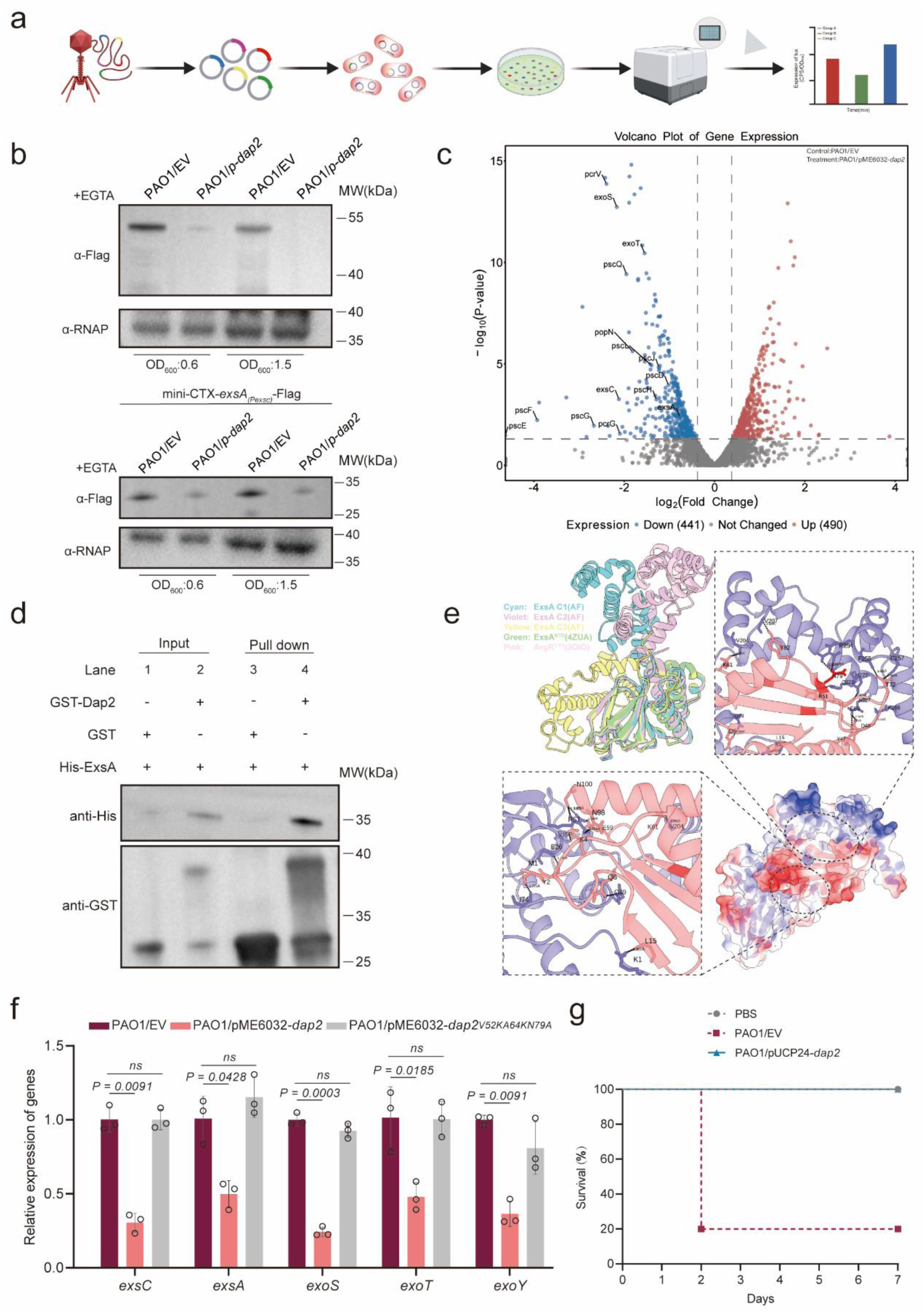
Phage protein Dap2 inhibits bacterial virulence via interacting with T3SS regulator ExsA. (a) Screening strategy for phage-encoded T3SS inhibitors. Schematic of the dual-plasmid screening system in *P. aeruginosa* PAO1. Early-expressed PaoP5 phage genes were cloned into IPTG-inducible plasmid pME6032. A T3SS activity reporter plasmid was constructed by fusing the *exsC* promoter to *luxCDABE* biosensor genes. Subsequently, the luminescence of each clone was measured using the microplate reader. (b) Western blot analysis of ExoS and ExsA protein expression in wild-type and Dap2-overexpressing PAO1 strains. Bacterial cultures were induced with 5 mM EGTA and 20 mM MgCl₂, with samples harvested at optical densities (OD₆₀₀) of 0.6 and 1.5. Protein levels were assessed using anti-ExoS and anti-Flag antibodies for ExoS and ExsA detection, respectively. α-RNA polymerase (RNAP) served as the loading control. (c) Volcano plot analysis of differentially expressed genes (DEGs) was performed using RNA-seq data comparing PAO1/p-*dap2* and PAO1/pME strains. Blue and red data points indicate genes with significant upregulation and downregulation, respectively, in PAO1/p-*dap2* relative to the PAO1/pME control group. (d) Direct interaction between GST-Dap2 and ExsA-His_6_ was confirmed by GST pull-down assay. Purified ExsA-His protein was incubated with either GST-Dap2 or GST (control) protein, and the resulting protein complexes were captured using GST-binding beads. The initial samples (input) and retained proteins (pull-down) were then analyzed by Western blot against GST or His antibody. (e) ExsA-Dap2 complex structure model and the details about interaction interface. Electrostatic potential surfaces of ExsA and Dap2 are basically complementary (bottom right). Different conformations of ExsA were marked with C1 to C3 (upper left). (f) Expression of *dap2* triple mutant (pME-*dap2*^V52K/A64K/N79A^) failed to inhibit T3SS-related genes. qRT-PCR analysis of T3SS gene expression in the indicated strains. Error bars represent the mean ± SD (n=3). Significant differences were assessed by an unpaired *t* test. ns, no significant difference. (g) The expression of *dap2* significantly attenuated the virulence of *P. aeruginosa*. In the infection model, 7-week-old BABL/c female mice were intranasally inoculated with either PAO1/pUCP24 (vector control) or PAO1/pUCP-dap2 at a dose of 2×10^7^ CFU in 100 μL of phosphate-buffered saline (PBS). Survival data were recorded. Statistical analysis using the Kaplan-Meier method (log-rank test) revealed a highly significant difference between the two groups (\*\*\*\**P* < 0.0001).

### Dap2 inhibits T3SS via interacting with ExsA

Given that ExsA serves as the master transcriptional regulator of T3SS, and our transcriptomic data revealed global downregulation of T3SS genes by Dap2 overexpression, we posited that Dap2 might directly target ExsA. To test this hypothesis, we first employed the BacterioMatch II Two-Hybrid System to probe Dap2-ExsA interactions. Recombinant strains expressing bait (pBT-Dap2) and prey (pTRG-ExsA) constructs were plated on nonselective versus dual-selective media containing 3-amino-1,2,4-triazole (3-AT) and streptomycin. While the positive control (LGF2/GAI11P interaction) showed robust growth under selection, specific colony survival was observed exclusively in the Dap2/ExsA co-expressing strain on selective media, demonstrating direct protein interaction (Fig. S3a). This physical interaction was further corroborated by affinity pull-down assays, where His-tagged ExsA co-purified with Dap2 (Fig. 1d). As anticipated, expression of *dap2* failed to repress *exoS* promoter activity in the Δ*exsA* mutant strain (Fig. S3b). Collectively, these results establish that Dap2-mediated suppression of T3SS occurs through direct binding to ExsA, thereby inhibiting its regulatory function.

Given ExsA’s role as a transcriptional activator of T3SS through promoter binding, we purified ExsA and Dap2 proteins (Fig. S3c) and conducted electrophoretic mobility shift assays (EMSAs) to determine whether Dap2 modulates ExsA-DNA interactions. EMSA revealed strong ExsA-dependent DNA binding, evidenced by a retarded band corresponding to ExsA-DNA complexes. Neither Dap2 nor the GST control protein exhibited DNA-binding activity (Fig. S3d). Strikingly, pre-incubation of ExsA with increasing Dap2 concentrations progressively attenuated complex formation, with 1 µM Dap2 completely abolishing the ExsA-DNA interaction (Fig. S3d). These results demonstrate that Dap2 interacts with ExsA to prevent its binding to DNA.

To analyze the potential interaction mode and identify key residues responsible for maintaining the ExsA-Dap2 interaction interface, molecular docking was conducted. Due to the existence of a hinge composed of the ^166’^Ser-Asn-Arg^168’^ triad, ExsA exhibits several conformations across all candidates. And it’s inferred that the relative motion ability of CTD (HTH domain) might contribute to the DNA binding. However, the domains located at each terminus are highly conserved compared to the reported ExsA^NTD^ or its homolog ArgR^CTD29^ (Fig. 1e, upper left). Therefore, different conformations of ExsA were docked with Dap2, whose structure was relatively conserved throughout all top candidates. The top ExsA-Dap2 model features an interface area of 2187.5 Å², accompanied by a solvation free energy gain of -13.7 kcal/mol, indicating a relatively hydrophobic interface or positive protein affinity as indicated by PDBePISA analysis. Besides 17 hydrogen bonds and 3 salt bridges, over 400 non-bonded contacts can be observed at the interface, suggesting that hydrophobic van der Waals forces might play a crucial role in maintaining the interface (Fig. 1e, bottom right). To validate the predicted interaction model, we generated site-directed mutants targeting key residues (V52K, A64K, N79A) and assessed their impact on T3SS gene promoter activity. Unlike wild-type Dap2, the triple mutant *dap2*^V52K/A64K/N79A^ failed to suppress T3SS genes (Fig. 1f). This functional impairment was corroborated by EMSAs, which revealed that both individual (V52K, A64K, N79A) and triple mutants lost the ability to disrupt ExsA-promoter complex formation (Fig. S3d-e). Notably, the control mutant E97A/E99A maintained wild-type levels of DNA binding suppression (Fig. S3f). These collective findings support a model where Dap2 interacts with ExsA to constrain conformational flexibility in its DNA-binding domain, thereby inhibiting ExsA-mediated transcriptional activation.

Given the critical role of the T3SS in virulence, we investigated the impact of the phage-encoded Dap2 protein on the pathogenicity of *P. aeruginosa* PAO1 using a well-established acute-infection mouse model. The *dap2* gene was cloned into the plasmid pUCP24 to enable constitutive expression of Dap2. The virulence of strains carrying either PAO1/pUCP24 (control) or PAO1/pUCP-*dap2* was then evaluated. Female BALB/c mice were intraperitoneally challenged with these strains, and survival was monitored. Strikingly, 80% of mice infected with PAO1/pUCP24 died within 2 days, whereas all mice infected with PAO1/pUCP-*dap2* survived through day 7 (Fig. 1g). Each experimental group included ten mice, and statistical analysis was performed using the Kaplan-Meier method (log-rank test). These findings demonstrate that the expression of Dap2 significantly attenuates the pathogenicity of *P. aeruginosa*.

### Knockout of *dap2* significantly impairs the fitness of phage

Our findings highlight the significant influence of the phage-encoded Dap2 protein on *P. aeruginosa* T3SS. To further explore the potential role of Dap2 in enhancing phage fitness, we investigated its transcriptional dynamics during infection. Specifically, *P. aeruginosa* strain PAO1 was infected with phage PaoP5, and samples were collected at 1-, 10-, 15-, and 30-minutes post-infection. RNA was extracted from these samples, and the expression levels of *dap2* transcripts were quantified using qRT-PCR. The results revealed that *dap2* transcription peaked at 10 minutes post-infection, while the mRNA level of structural gene *orf053* was highest at 30 minutes (Fig. S4a). This temporal expression pattern indicates that *dap2* is an early-expressed gene, likely to play a role in the initial stages of phage infection.

Next, we employed the CRISPR-Cas9 system to delete the *dap2* gene in phage PaoP5, with the knockout of *orf014* serving as a control. Notably, PaoP5Δ*dap2* produced significantly smaller plaques compared to the wild-type (WT) phage, while PaoP5Δ*orf014* formed plaques similar in size to the WT (Fig. 2a). Furthermore, the expression of *dap2* in *P. aeruginosa* PAO1 restored the formation of large plaques for PaoP5Δ*dap2*, whereas the efficiency of plating (EOP) of PaoP5Δ*dap2* in PAO1/p-dap1 remained comparable to that in PAO1/EV (Fig. 2b). These results demonstrate that *dap2* plays a critical role in phage fitness, whereas *orf014* does not appear to contribute to phage fitness under the tested laboratory conditions.

**Fig. 2.**
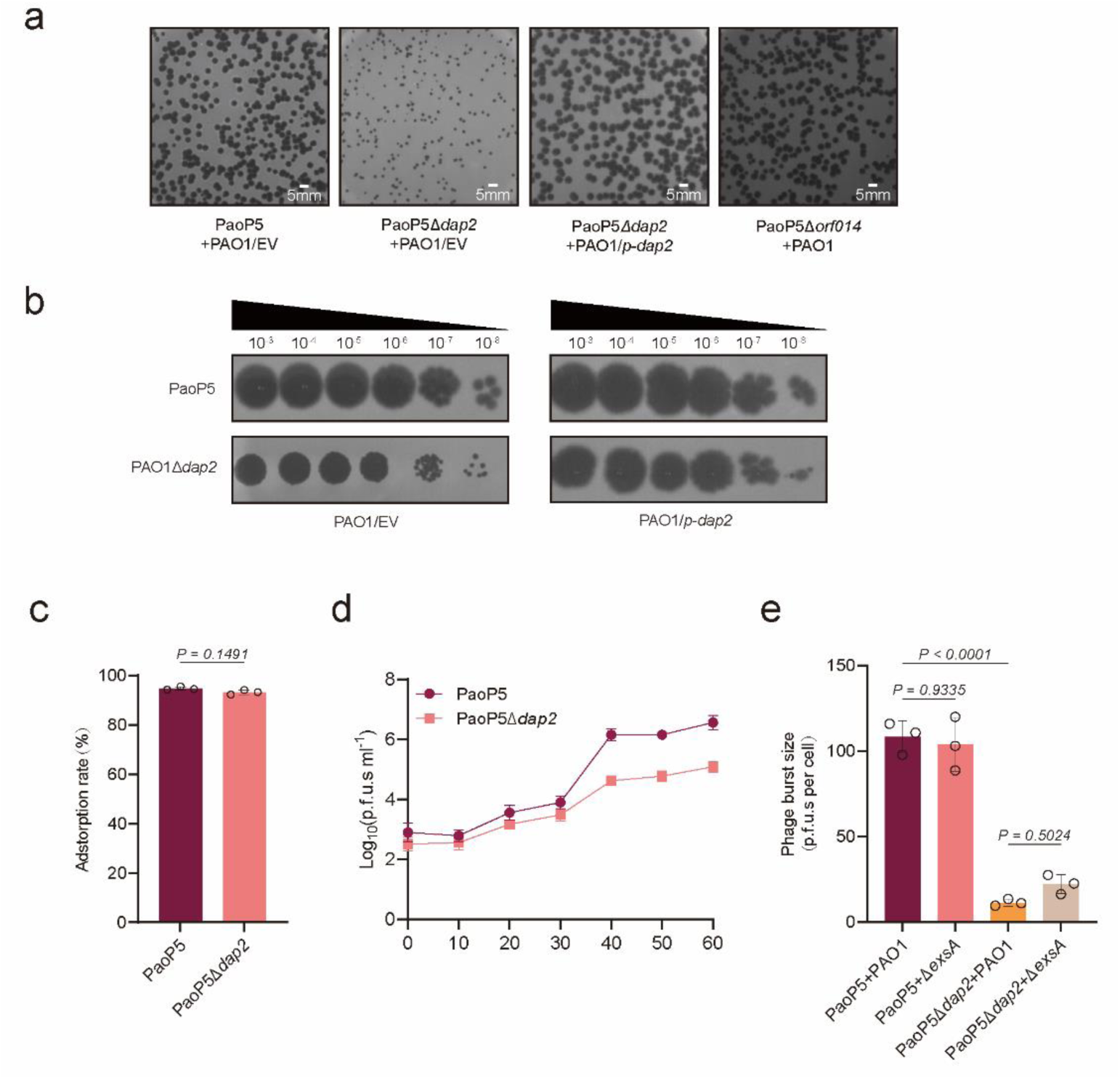
PaoP5Δ*dap2* forms small plaques and produces fewer progenies. (a) PaoP5*Δdap2* produced significantly smaller phage plaques compared to the wild-type PaoP5, while PaoP5*Δorf014*, used as a control, exhibited plaque sizes similar to the wild-type. Complementation with Dap2 restored the large plaque phenotype, indicating the functional role of Dap2 in plaque formation. (b) Plaque assays were performed using 10-fold serial dilutions to compare the plating efficiency of wild-type (WT) and mutant phages on PAO1 and PAO1 harboring the *p-dap2* vector. The results demonstrate the impact of Dap2 on phage plaque size but do not affect EOP. (c) Both PaoP5Δ*dap2* and wild-type PaoP5 phages adsorbed to PAO1 with comparable efficiency. Error bars represent the mean ± SD of four biological replicates. Statistical analysis using Student’s t-test showed no significant difference (ns, not significant). (d) The one-step growth curve revealed that PaoP5*Δdap2* produced significantly fewer progeny phages compared to the wild-type PaoP5, highlighting the role of Dap2 in phage replication efficiency. (e) The burst size of wild-type PaoP5 infecting PAO1 and PAO1Δ*exsA* was 1108.26± 6.65 PFU/cell and 103.91±11.08 PFU/cell, respectively. In contrast, the burst size of PaoP5Δ*dap2* infecting PAO1 and PAO1Δ*exsA* was significantly reduced to 11.32±1.42 PFU/cell and 22.08±3.99 PFU/cell, respectively. Error bars represent the mean ± SD of three biological replicates. Statistical significance was determined using Student’s *t*-test.

To investigate the role of *dap2* in the phage life cycle, we first assessed its impact on the initial adsorption rate, as phage infection begins with host recognition and binding. Both PaoP5Δ*dap2* and the wild-type (WT) phage exhibited similar adsorption efficiencies, with no statistically significant difference (*P* > 0.05), indicating that *dap2* is not involved in host recognition or binding (Fig. 2c). We then examined the burst size of the two phages, as plaque size is closely linked to the number of progeny produced^30^. The one-step growth curve revealed that PaoP5Δ*dap2* generated significantly fewer progeny compared to the WT phage (Fig. 2d). Specifically, the burst sizes of PaoP5 and PaoP5Δ*dap2* were approximately 108.26 ± 6.65 PFU/cell and 11.32 ± 1.42 PFU/cell, respectively (Fig. 2e). This indicates that PaoP5Δ*dap2* produced only about 10.45% of the progeny generated by the WT phage.

To determine whether the reduced productivity was linked to the bacterial T3SS, we infected PAO1Δ*exsA* with PaoP5Δ*dap2*. However, the small plaque phenotype persisted (Fig. S4c), suggesting that the fitness advantage conferred by Dap2 extends beyond its interaction with ExsA.

Since larger plaques require a high yield of progeny per generation^30^, subtle differences in burst size may not be discernible through plaque morphology alone. To further explore this, we infected PAO1 and PAO1Δ*exsA* with PaoP5Δ*dap2*. Interestingly, the burst size of PaoP5 infecting PAO1 or PAO1Δ*exsA are* 108.26 ± 6.65 PFU/cel and 103.91±11.08 PFU/cel, respectively, which is not significantly different (*P*=0.7014), the burst size of PaoP5Δ*dap2* in PAO1Δ*exsA* was 22.08±3.99 PFU/cell compared to 11.32±1.42 PFU/cell in *PAO1* (*P*=0.0357) (Fig. 2e), indicating that the phage’s inhibition of T3SS serves as a specific strategy to reduce host energy consumption, thereby enhancing progeny production. This provides direct evidence of the fitness benefits conferred by Dap2 through T3SS suppression. However, the small plaque phenotype of PaoP5Δ*dap2* was not reversed in PAO1Δ*exsA*, suggesting that Dap2 may also function as an anti-defense system, targeting an additional bacterial defense mechanism beyond T3SS. This dual role highlights the multifaceted nature of Dap2 in promoting phage fitness.

### Dap2 binds to Lon protease to inhibit its degradation on the phage HNH endonuclease

To identify the potential anti-defense target of Dap2, we employed a pull-down assay coupled with liquid chromatography-mass spectrometry (LC-MS) to screen for Dap2-interacting proteins. This analysis identified Lon as a binding partner of Dap2 (Fig. 3a, Table S4). The interaction between Dap2 and Lon was further validated using a bacterial adenylate cyclase two-hybrid (BACTCH) assay (Fig. 3b), which confirmed their specific binding. Notably, Dap2 did not interact with Dap1 or HNH under the same conditions. The Dap2-Lon interaction was additionally confirmed through a pull-down assay, reinforcing the specificity of this binding (Fig. 3c).

**Fig. 3.**
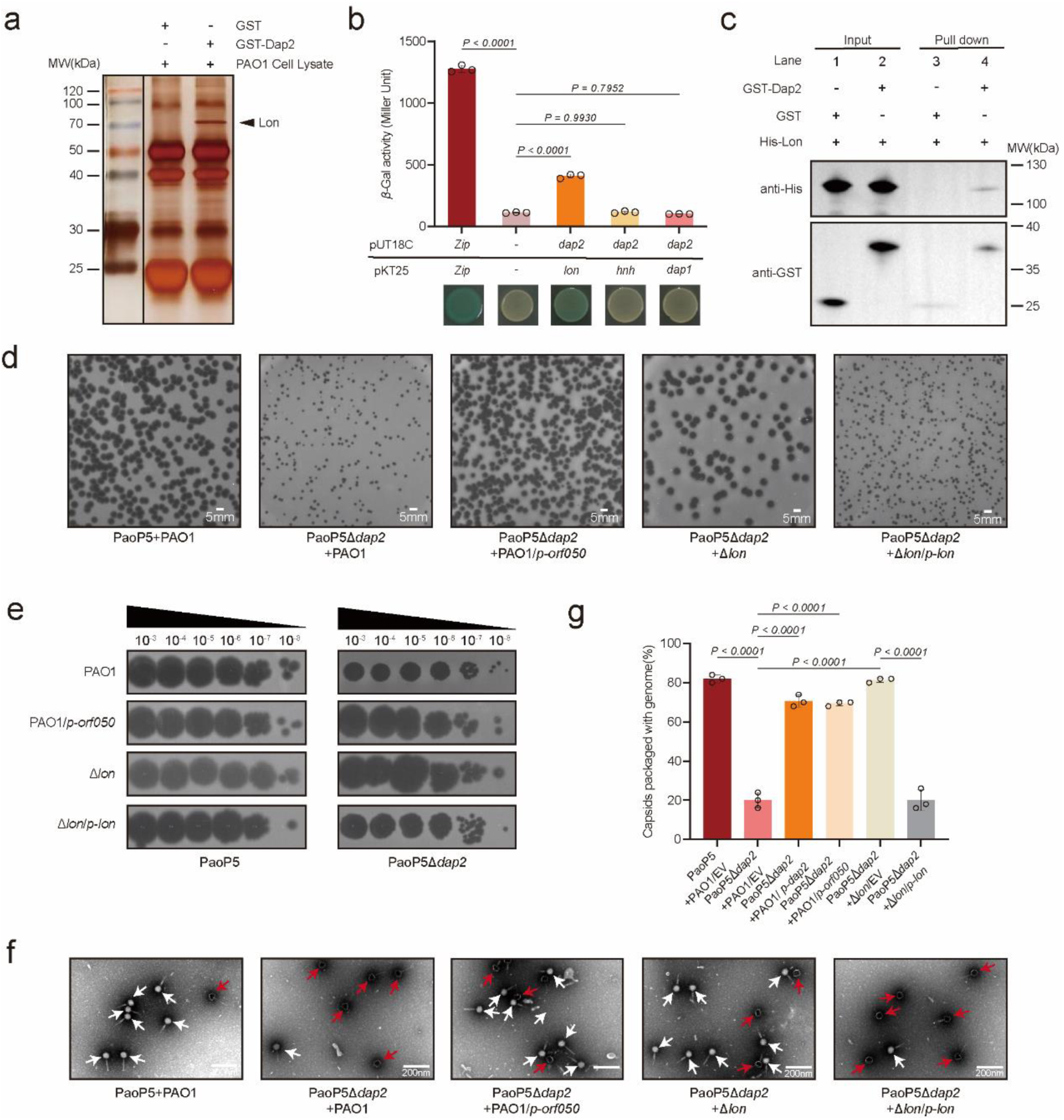
Dap2 binds to Lon protease to inhibit its function. (a) Identification of Lon as the binding target of Dap2. Proteins specifically retained by GST-Dap2 from cell lysates of P. aeruginosa PAO1 with a silver-stained SDS–PAGE gel. The identity of the retained protein (red arrow) was identified through mass spectrometry. (b) The bacterial two-hybrid assay revealed that Dap2 specifically interacts with Lon protease, but not with HNH or Dap1. These interactions were visualized through a drop test on LB agar plates, where the formation of blue colonies signified positive interactions. The interactions were further quantified by measuring β-galactosidase activity. Error bars represent the mean±SD of three biological replicates. Statistical significance was determined using one-way ANOVA with Dunnett’s multiple comparison test: ns, not significance. (c) Direct interaction between GST-Dap2 and Lon-His_6_ was confirmed by GST pull-down assay. Purified Lon-His protein was incubated with either GST-Dap2 or GST (control) protein, and the resulting protein complexes were captured using GST-binding beads. The initial samples (input) and retained proteins (pull-down) were then analyzed by Western blot against GST or His antibody. (d) Plaque formation and EOP (e) was observed for PaoP5 infecting PAO1, as well as PaoP5Δ*dap2* infecting PAO1*/p-dap2*, Δ*lon*, Δ*lon/p-lon*, and PAO1/*p-hnh*. (f) Transmission electron microscopy (TEM) images of negatively stained phages produced in PAO1, Δ*lon,* Δ*lon/p-lon*, and PAO1/*p-hnh* are shown. Red arrows indicate empty capsids, while white arrows point to phages with packaged genomes. (g) The percentage of capsids containing genomes was calculated from three biological replicates. For each condition, 50 particles were counted to determine the presence or absence of genomes. Error bars represent the mean ± SD of three biological replicates. Statistical significance was determined using one-way ANOVA with Dunnett’s multiple comparison test.

Previously, we demonstrated that Lon protease directly degrades the phage-encoded HNH endonuclease^22^, a critical component of the phage DNA packaging machinery required for the specific endonuclease activity of large terminase proteins^26^. Degradation of HNH disrupts phage genome packaging, suggesting that Dap2 may bind to Lon to inhibit its function. To test this hypothesis, we infected wild-type PAO1, *Δlon*, and *Δlon/p-lon* strains with PaoP5Δ*dap2*. The results showed that PaoP5Δ*dap2* formed larger plaques in *Δlon* or PAO1*/p-hnh* compared to PAO1 (Fig. 3d-e), while plaque size reverted to smaller dimensions in *Δlon/p-lon*, supporting the role of Lon in HNH degradation.

Since Lon-mediated degradation of HNH leads to the formation of empty capsids devoid of packaged phage genomes, we examined phage lysates from different strains using transmission electron microscopy (TEM). Negatively stained electron micrographs revealed a significant number of empty phage particles in PaoP5Δ*dap2* lysates, indicative of defective DNA packaging. In contrast, deletion of *lon* or overexpression of *hnh* in PAO1 restored DNA packaging efficiency to levels comparable to the wild-type phage (Fig. 3f, Fig. S7).

Quantification of genome-packaged capsids from three biological replicates showed that only 20.00% ± 3.26% of capsids contained genomes in PaoP5Δ*dap2* lysates. However, complementation with *dap2*, overexpression of *hnh*, or deletion of *lon* increased this percentage to 70.67%±2.49%, 69.33%±0.94%, and 81.33% ± 0.94%, respectively. Conversely, reintroduction of *lon* into *Δlon* decreased the percentage of genome-packaged capsids to 20.00 ± 4.32% (Fig. 3g). These findings demonstrate that Dap2 protects HNH endonuclease from rapid degradation by Lon, thereby ensuring efficient phage genome packaging and maintaining the productivity of phage progeny.

### Dap2 and Dap1 cooperate to evade the Lon protease-mediated anti-phage defense

Building on our prior finding that Dap1 sterically shields the HNH endonuclease from Lon protease-mediated degradation^22^, we investigated potential functional synergy between Dap1 and Dap2. The genomic colocalization of these adjacent genes within a conserved operon, coupled with Dap2’s direct Lon-binding capacity, prompted us to propose a cooperative defense mechanism ensuring full protection of HNH.

To confirm this hypothesis, we first generated a double knockout of *dap1* and *dap2*. The resulting phage, PaoP5Δ*dap1*Δ*dap2*, formed tiny, blurred plaques, significantly smaller than those produced by PaoP5Δ*dap1* or PaoP5Δ*dap2* alone (Fig. 4a). TEM analysis revealed that ∼90% of the capsids were empty (Fig. 4b), indicating severe defects in genome packaging. Complementation of HNH or deletion of Lon in PAO1 could restore the large plaque size (Fig. S6a) and increase the genome packaging efficiency (Fig. S6b). While adsorption kinetics remained unaffected in the double mutant (Fig. S6c), the one-step growth curve demonstrated a delayed burst time and a markedly reduced burst size for PaoP5 *Δ dap1 Δ dap2* (Fig. 4c). The burst size decreased to 7.09±0.37 PFU/cell, only ∼6.68% of that observed for the wild-type phage (Fig.S6d), highlighting the synergistic role of Dap1 and Dap2 in countering Lon-mediated defense.

**Fig. 4.**
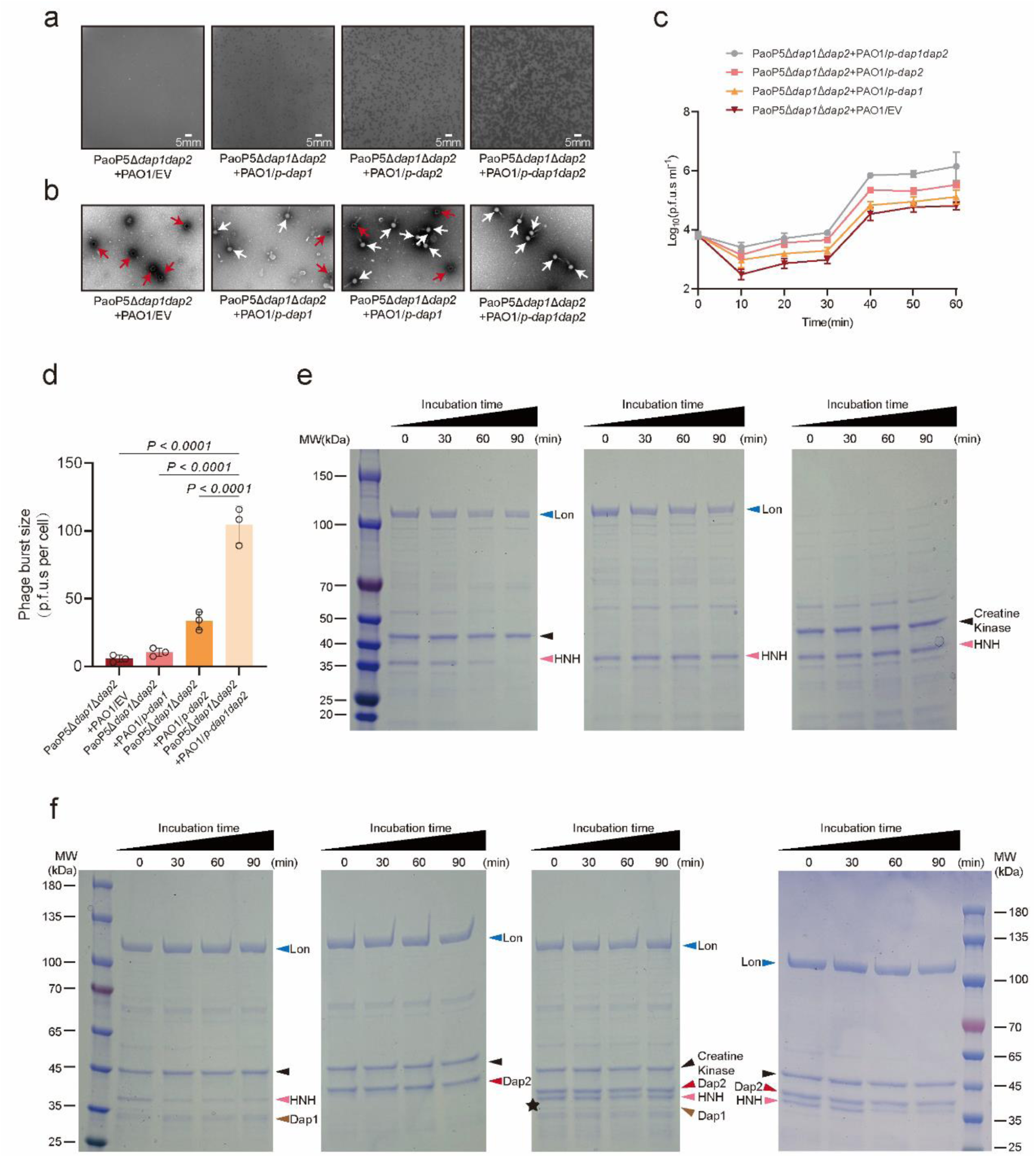
Dap2 and Dap1 cooperate to defend Lon protease. (a) Plaque formation of PaoP5*Δdap1Δdap2* was assessed on PAO1, PAO1/*p-dap1*, PAO1/*p-dap2*, and PAO1/*p-dap1dap2*, demonstrating the impact of Dap1 and Dap2 complementation on phage infectivity. (b) Transmission electron microscopy (TEM) images of negatively stained PaoP5Δ *dap1*Δ*dap2* phages produced in PAO1, PAO1/*p-dap1*, PAO1/*p-dap2*, and PAO1/*p-dap1dap2* are shown. Red arrows highlight empty capsids, while white arrows indicate phages with packaged genomes. (c) The one-step growth curve revealed that PaoP5 Δ *dap1* Δ*dap2* produced fewer progeny phages in PAO1 compared to the wild-type. However, complementation with both Dap1 and Dap2 significantly restored the burst size, underscoring their collective role in phage replication. (d) The burst size of PaoP5*Δdap1Δdap2* infecting PAO1, PAO1/*p-dap1*, PAO1*/p-dap2*, and PAO1*/p-dap1dap2* was 5.66 ± 1.88 PFU/cell, 10.44 ± 2.11 PFU/cell, 33.66 ± 4.73 PFU/cell, and 104.46 ± 9.76 PFU/cell, respectively. Error bars represent the mean ± SD of three biological replicates. Statistical significance was determined using Student’s *t*-test. (e-f) *In vitro* proteolysis assays demonstrated that the combination of Dap1 and Dap2 completely prevented Lon-mediated degradation of HNH, whereas either protein alone only slowed the process. Complete substrate stabilization was achieved throughout the 90-minute reaction when both proteins were present. Protein samples collected at specified time points were analyzed by 12% SDS–PAGE and visualized via Coomassie staining. All experiments were repeated at least three times with consistent results, and representative data are shown.

To further validate this synergy, we performed complementation assays. Individual *dap1* or *dap2* expression partially rescued plaque morphology, genome packaging, and burst size, whereas co-expression fully restored wild-type parameters (Fig. 4a-d). We also conducted *in vitro* protein degradation assays using purified Lon, HNH, ATP, and an ATP regeneration system (creatine phosphate and creatine phosphokinase)^22,31^. HNH was completely degraded in the presence of Lon and the kinase system but remained stable in the absence of either creatine phosphokinase or Lon (Fig. 4e). Intriguingly, Lon was unable to degrade Dap2, and Dap2 alone provided partial protection against HNH degradation. Similarly, we previously showed that Dap1 also offers partial protection to HNH^22^. Remarkably, when Dap1 and Dap2 were combined, they completely inhibited LON-mediated degradation of HNH (Fig. 4f).

These *in vitro* and *in vivo* findings demonstrate that Dap1 and Dap2 form a synergistic anti-defense system (ADS) pair, employing distinct mechanisms to neutralize Lon protease: Dap1 shields the target (HNH), while Dap2 directly inhibits the defense protein (Lon). Individually, each protein provides only partial protection, but together, they achieve complete protection, highlighting the evolutionary advantage of this dual-defense strategy.

### Dap2/Dap1 Pair is Co-localized in Phage Genomes and Cooperates to Enhance the Efficacy of Phage Therapy

To explore the prevalence of the *dap1/dap2* gene pair, we conducted a homology search in the NCBI database among *P. aeruginosa* phages. As of January 2025, 67 sequenced *Pseudomonas* phages were found to carry *dap2* homologs with a minimal identity of 80%, which were clustered into three distinct clades based on phylogenetic analysis (Fig. 5a).

**Fig. 5.**
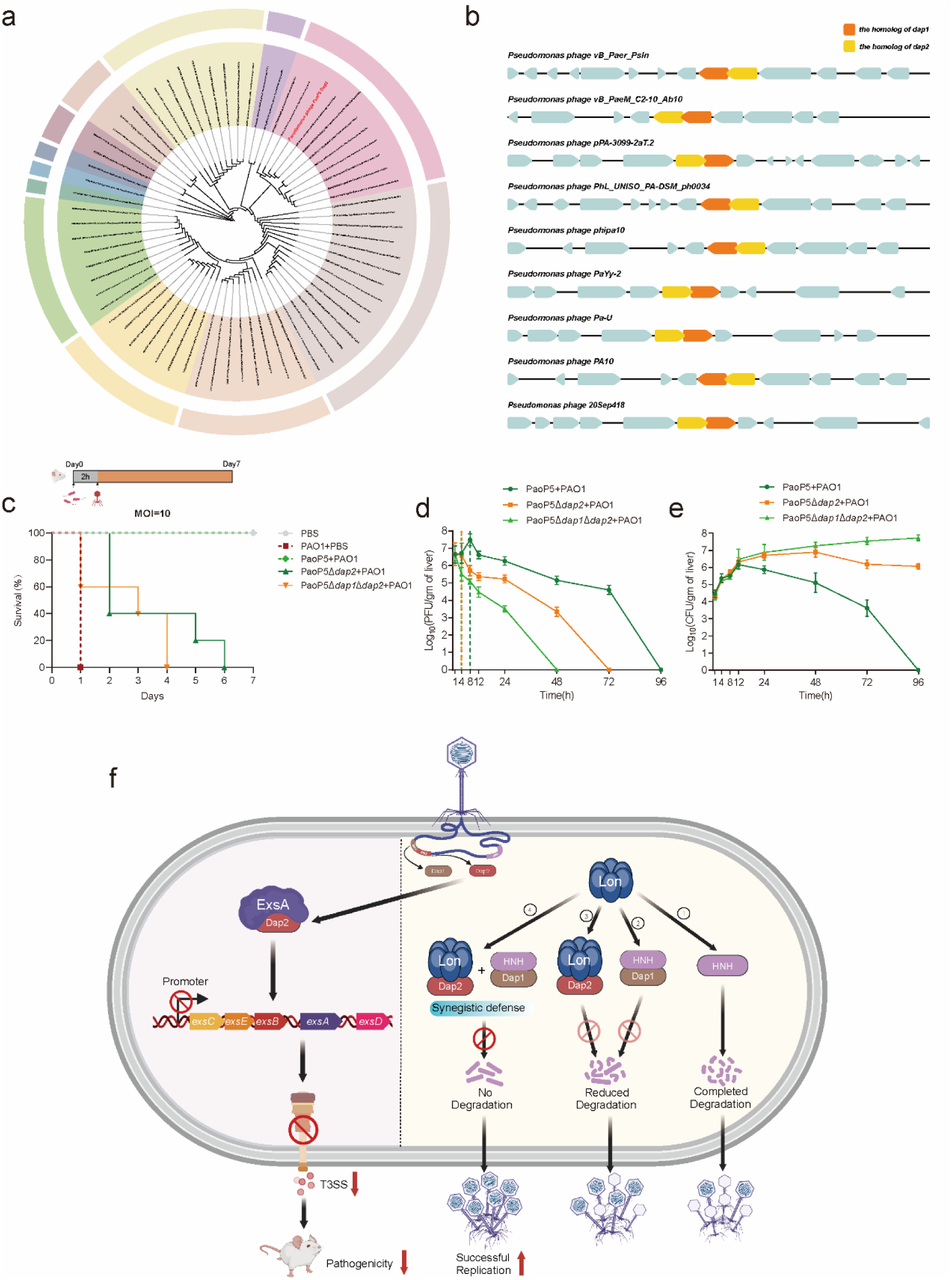
Dap2/Dap1 Pair is Co-localized in Phage Genomes and Cooperates to Enhance the Efficacy of Phage Therapy. (a) A phylogenetic tree of Dap2 homologues with a minimum identity of 80% was constructed. The tree was generated using neighbor-joining analysis in MEGA 7, with evolutionary distances calculated using the p-distance method. The phylogenetic trees were visualized using iTOL. (b) Dap1 and Dap2 homologues, which are tandem overlapping genes, were identified in 67 phage genomes using BLAST. Nine representative phage genomes containing Dap1 and Dap2 homologues are illustrated. (c) Survival of 7-week-old BALB/c mice (n = 5) following intraperitoneal injection with PAO1 and phage PaoP5Δ*dap2*, PaoP5Δ*dap1*Δ*dap2*, or PaoP5 at an MOI of 10. (d) Phage titers and (e) bacterial CFUs were measured in the livers of mice intraperitoneally injected with PAO1 and phage PaoP5Δ*dap2*, PaoP5Δ*dap1*Δ*dap2*, or PaoP5 at an MOI of 10. The maximum titer of PaoP5 was detected at 8 hours post-injection (∼3.8 × 10^7^ PFU/g). The phage titer gradually decreased after 24 hours and was cleared within 96 hours. Concurrently, bacterial CFUs in the liver decreased steadily and were fully cleared within 96 hours. In contrast, for PaoP5Δ*dap1*Δ*dap2-* treated, PAO1-infected mice, the phage titer continuously decreased and became undetectable by 48 hours, with a maximum titer of only ∼8.25 ×10^6^ PFU/g-significantly lower than that observed in PaoP5-treated mice. Error bars represent the mean ± standard deviation (n = 4 mice). (f) Proposed model depicting the dual mechanisms by which Dap2 suppresses bacterial T3SS and collaborates with Dap1 to counteract Lon-mediated anti-phage defense. The phage-encoded protein Dap2 interacts with ExsA, the master regulator of the T3SS, thereby inhibiting T3SS expression and significantly attenuating the virulence of *P. aeruginosa*. Furthermore, during PaoP5 infection of PAO1, Dap1 binds to the HNH endonuclease, protecting it from Lon-mediated degradation, while Dap2 directly interacts with the Lon protease, effectively inhibiting its activity. Individually, Dap1 or Dap2 offers only partial protection against Lon-mediated anti-phage defense. However, their combined action completely neutralizes Lon protease activity, ensuring the preservation of the HNH endonuclease and facilitating efficient phage genome packaging. This synergistic interplay between Dap1 and Dap2 guarantees the production of sufficient phage progeny, exemplifying a refined evolutionary adaptation to overcome host immune defenses.

Interestingly, *dap1* and *dap2* genes consistently co-occur, forming a tandem overlapping gene pair separated by *a* base pair (Fig. 5b). This gene pair was identified in 67 phage genomes (Table S5), suggesting a conserved genetic arrangement. This widespread co-localization indicates that *dap1* and *dap2* function as a synergistic anti-defense system (ADS) pair, both essential for countering the Lon protease, a ubiquitous housekeeping protein in *P. aeruginosa*. The evolutionary conservation of this gene pair underscores its critical role in enhancing phage fitness and efficacy during infection.

Given that phage PaoP5*Δdap2* produces fewer progeny and PaoP5*Δdap1Δdap2* generates even fewer, we assessed their therapeutic potential using a mouse infection model. Seven-week-old female BALB/c mice were intraperitoneally injected with either 50 μL of PBS (uninfected control) or 50 μL of *P. aeruginosa* PAO1 (3 × 10^8^ CFU/mL). This was followed by administration of 50 μL of phage PaoP5*Δdap1Δdap2,* PaoP5*Δdap2,* or wild-type PaoP5 at a multiplicity of infection (MOI) of 10. Wild-type PaoP5 rescued 100% of *P. aeruginosa*-infected mice (Figure 6b). In contrast, all mice treated with PaoP5*Δdap2* succumbed to infection within 6 days, while those treated with PaoP5*Δdap1Δdap2* died even more rapidly, with all mice perishing within 4 days (Fig. 5c).

To further evaluate the therapeutic efficacy of PaoP5Δ*dap2* and PaoP5Δ*dap1*Δ*dap2*, we measured phage titers and bacterial CFUs in the liver post-treatment. As shown in Fig. 5d, significantly higher phage titers were detected in the livers of PaoP5-treated, PAO1-infected mice. The peak titer of PaoP5 reached ∼3.8 × 10^7^ PFU/g at 8 hours post-injection, gradually declining thereafter and clearing within 96 hours (Figure 5d). Concurrently, bacterial CFUs in the liver decreased steadily and were fully cleared by 96 hours, resulting in 100% survival of PaoP5-treated mice (Fig. 5e). In contrast, for PaoP5*Δdap2*-treated, PAO1-infected mice, phage titers continuously decreased, becoming undetectable by 72 hours, with a maximum titer of only ∼1.1 × 10^7^ PFU/g-significantly lower than that observed in PaoP5-treated mice. Similarly, PaoP5*Δdap1Δdap2*-treated, PAO1-infected mice exhibited even lower phage titers during the therapy. Bacterial CFUs in the liver continued to rise post-infection, ultimately leading to the death of all PaoP5*Δdap1Δdap2*- and PaoP5*Δdap2*-treated mice. These results demonstrate that PaoP5*Δdap1Δdap2* produces even fewer progeny, significantly impairing its therapeutic efficacy. Collectively, these findings highlight the critical role of both *dap1* and *dap2* in the effectiveness of phage therapy.

## Discussion

The relentless evolutionary arms race between bacteriophages and their bacterial hosts has driven the diversification of phage-encoded anti-defense systems (ADSs) to overcome host immunity^10^. While ADSs targeting established bacterial defenses like CRISPR-Cas and restriction-modification (R-M) systems are well characterized^32^, recent discoveries have unveiled ADSs countering emerging immune pathways such as CBASS, Pycsar, Gabija, Thoeris, and Hachiman^11,13^. Despite these advances, critical gaps persist in ADS biology. For example, certain ADSs exhibit incomplete protective capacity, exemplified by the phage protein Acb2, which only partially rescues phage titers in JBD67*Δacb2* and JBD18 variants^16^. This partial efficacy highlights two key insights: (1) the need for quantitative assessments to define ADS protective thresholds, and (2) the potential existence of synergistic ADS pairs that act cooperatively to fully neutralize bacterial defenses^10^.

Our study reveals a novel cooperative ADS mechanism. Previously, we demonstrated that Dap1 binds to the phage-encoded HNH endonuclease, providing partial protection against Lon protease-mediated degradation^22^. Here, we show that Dap2 directly binds to and inhibits Lon’s proteolytic activity. *In vitro* reconstitution experiments demonstrated that combined expression of Dap1 and Dap2 completely blocks HNH degradation. *In vivo* validation in *P. aeruginosa* PAO1 strains revealed that low-concentration expression of either Dap1 or Dap2 only partially restored plaque formation in PaoP5*ΔDap1Δdap2* phages. Strikingly, co-expression of both proteins at low concentrations fully restored plaque morphology and increased burst size in PAO1::Dap1/Dap2 strains (Fig. 4a). Notably, the PaoP5Δ*Dap1*Δ*dap2* phage exhibited further reduced burst sizes due to HNH deficiency, underscoring the critical role of this ADS pair in phage fitness and therapeutic efficacy.

This synergy represents a compelling example of coordinated ADS action against Lon protease. Our findings suggest an evolutionary strategy where phages deploy complementary ADS combinations when single systems are insufficient. Genomic analysis revealed widespread conservation of *dap1/dap2*-like tandem overlapping gene pairs across phages (Fig. 5b), indicating their essential role in countering host immunity. This genomic colocalization further supports the hypothesis that synergistic ADS pairs are evolutionarily favored, and investigating genes adjacent to known ADS loci may uncover novel defense-countering systems^33^.

Beyond its anti-defense role, Dap2 exhibits dual functionality by suppressing bacterial virulence. We identified Dap2 as a direct binder of ExsA, the master regulator of *P. aeruginosa’s* T3SS^24^, leading to T3SS suppression and attenuated pathogenicity in a murine acute-infection model (Fig. 1h). This highlights a phage strategy to modulate host metabolism for survival advantage. While most studies focus on phage-encoded lethal proteins targeting transcription or replication^18,34^, our work emphasizes the untapped potential of virulence-modulating phage proteins as anti-infective agents^35^.To explore the biological benefit of T3SS inhibition, we compared phage progeny production in PAO1 and Δ*exsA* strains. While Δ*exsA* did not rescue the small plaque phenotype of PaoP5Δ*dap2*, burst size analysis revealed a ∼1.95-fold increase in progeny production when PaoP5Δ*dap2* infected *ΔexsA* compared to wild-type PAO1. Although this fitness gain appears modest, cumulative effects from multiple phage proteins targeting diverse host pathways could significantly enhance phage replication efficiency.

This study elucidates the dual functionality of a ADS protein Dap2, which simultaneously targets the virulence regulator ExsA and the Lon protease — two unrelated bacterial proteins—to maximize phage progeny production. Furthermore, we unveil a synergistic ADS pair (Dap1/Dap2) that employs distinct mechanisms to fully neutralize Lon-mediated defense, exemplifying a sophisticated evolutionary strategy. These findings advance our understanding of phage counter-defense tactics and highlight the potential of dual-function phage proteins as blueprints for novel antimicrobial and anti-virulence therapies.

## AUTHOR CONTRIBUTIONS

Conceptualization: H.L., and S.L

Methodology: J. Z., Y. Z., and C.

W Validation: H.L., and S. L

Investigation: J. Z., Y. Z., C. W., F. T., J.L. L., J. D., Z. Z., J. L., and N. G

Data curation: J. Z., Y. Z., and F. T

Writing – review & editing: J. Z., S. L., and H. L

## Acknowledgments

This study was supported by grants from the National Natural Science Foundation of China (32170188 to H. L), the National Key Research and Development Program of China (2022YFC2304401, 2021YFA0911200), the Shenzhen Science and Technology Program (20231120104808001), and Key University Laboratory of Metabolism and Health of Guangdong, SUST. The funders had no role in study design, data collection and interpretation, or the decision to submit the work for publication.

## Figure Legends

**Fig. S1.**
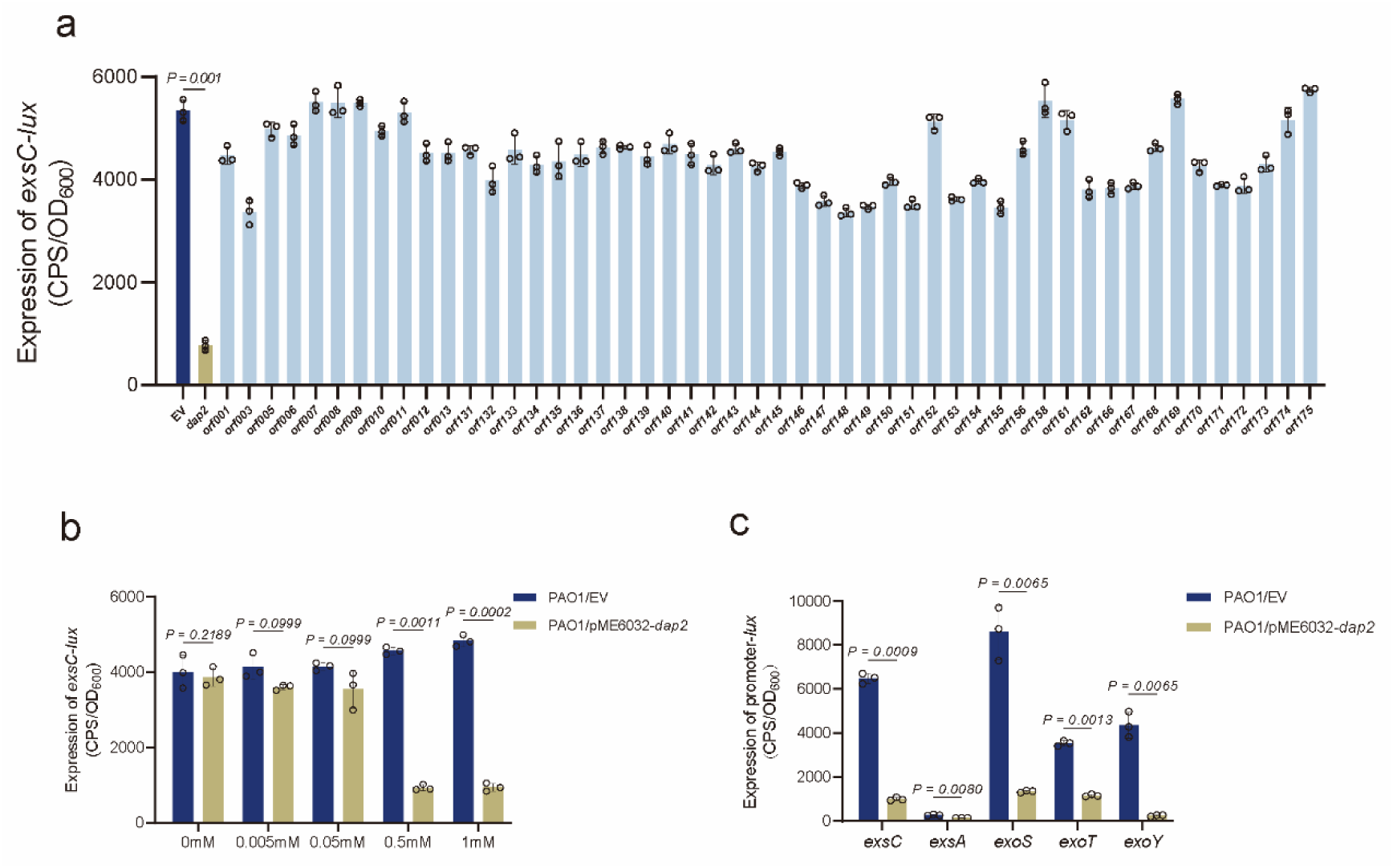
Overexpression of *dap2* inhibited the activity of T3SS-related genes. (a) The promoter activity of *exsC* was assessed in strains carrying either *exsC-lux* or 51 hypothetical ORFs derived from PaoP5. These constructs were individually expressed using the pME6032 vector and cultured in LB medium supplemented with 5 mM EGTA, 20 mM MgCl₂, and 0.05 mM IPTG. Promoter activity measurements were performed following 6 hours of cultivation (b) The expression of *exsC* in PAO1/pME6032 and PAO1/pME6032-*dap2* cultured in T3SS inducing medium supplemented with different concentrations of IPTG. (c) The promoter activities of pKD-*exsA*, pKD-*exsC*, pKD-*exoS*, pKD-*exoT*, and pKD-*exoY* were comparatively analyzed in PAO1/EV and PAO1-pME6032-*dap2* strains under T3SS-inducing conditions supplemented with 0.5 mM IPTG. Following 6-hour cultivation, transcriptional activity measurements revealed differential expression patterns. (a-c) Error bars indicate the mean ± SD (n=3) and statistical significance was determined using a two-sided Student’s *t-*test. EV, empty vector.

**Fig. S2.**
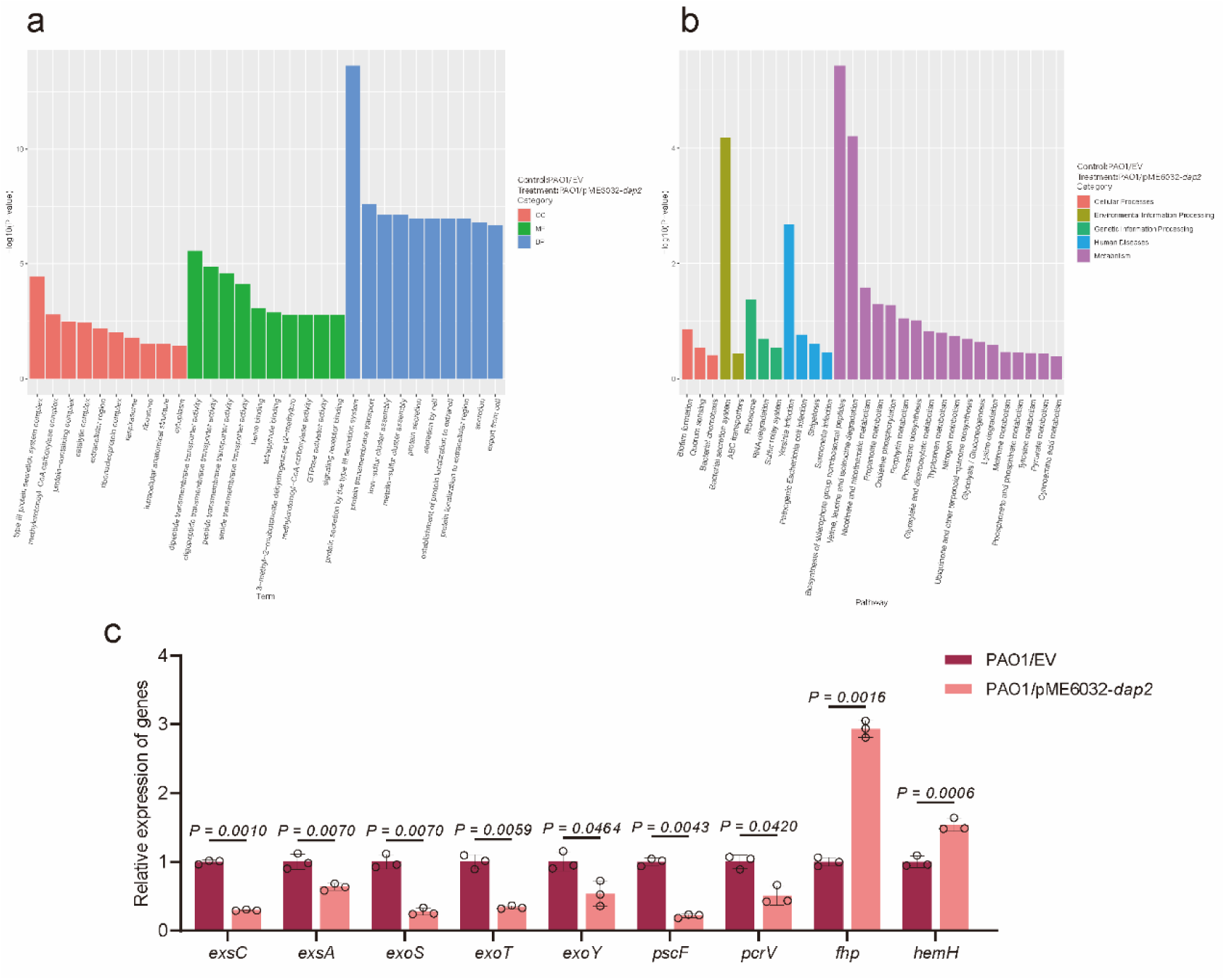
RNA-seq analysis between PAO1 and PAO1/pME6032-*dap2*. (a) The GO enrichment of the differentially expressed genes in PAO1/EV and PAO1/ pME6032*-dap2* strains, which is classified according to molecular function (MF), biological process (BP), and cellular component (CC), and the top 10 enriched GO was shown. (b) The DEGs are classified based on the KEGG analysis, and the top 30 enriched pathways, including Quorum Sensing and biofilm formation, are displayed. (c) Comparative qRT-PCR analysis of target gene expression in PAO1/EV versus PAO1/pME6032-*dap2* strains under T3SS-inducing conditions. Quantitative measurements from three biological replicates (mean ± SD, n = 3) demonstrated statistically significant variations (two-tailed Student’s t-test). EV, empty vector.

**Fig. S3.**
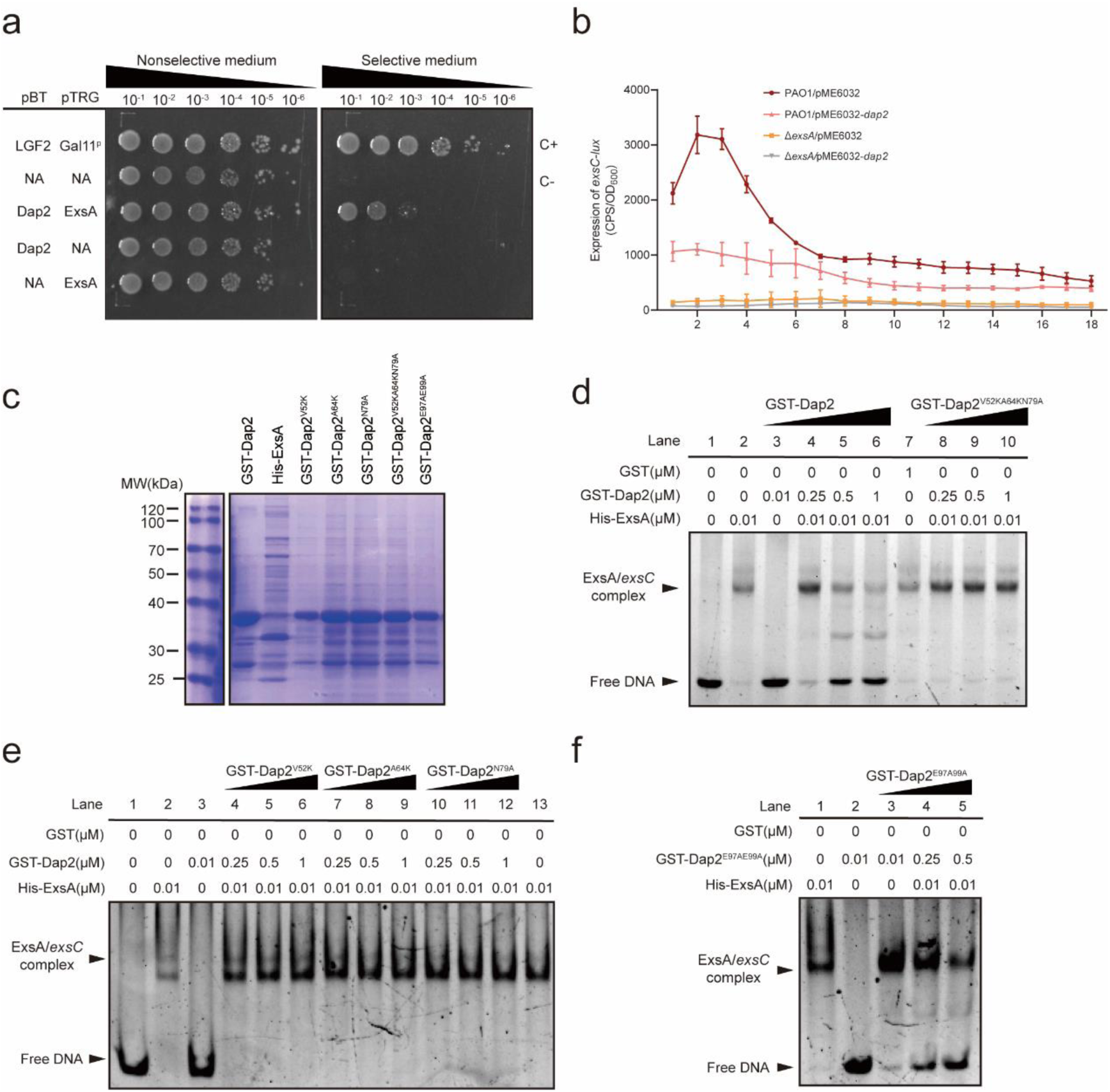
Verification of the interaction between Dap2 and ExsA *in vitro* and *in vivo*. (a) *E. coli* two-hybrid assay reveals an interaction between Dap2 and ExsA. The recombinant strains harboring different plasmids were separately streaked on nonselective and dual-selective media. The strain expressing LGF2 and GAI11P was used as a positive control. (b) Expression of *dap2* did not suppress *exsC* activity in Δ*exsA* mutant strain. *P. aeruginosa* carrying pKD-*exsC* with pME6032-*dap2* was cultivated in T3SS-inducing medium. The promoter activity was detected at specified time points. Error bars represent mean ± SD of three biological replicates. (c) Purified ExsA, Dap2 and Dap2-derived mutants were resolved on 12% SDS-polyacrylamide gels and stained with Coomassie Brilliant Blue R-250. (d-f) EMSA showing that Dap2 affects the binding ability of ExsA to DNA. A mixture of 10 ng DNA and His-ExsA was incubated with increasing concentrations of GST-Dap2 or GST-Dap2 mutant proteins. Data are representative of three independent replicates.

**Fig. S4.**
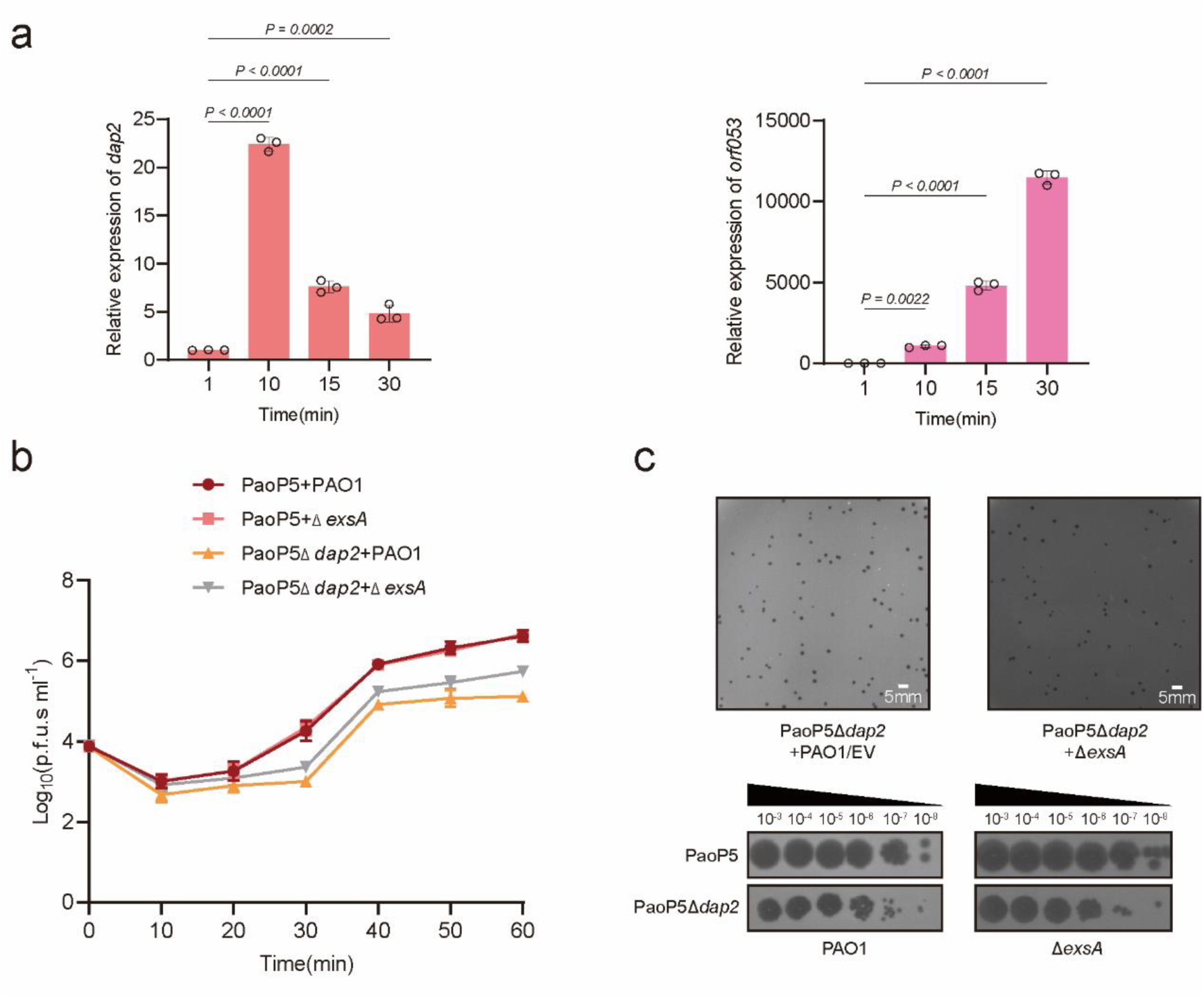
Dap2 is an early expressed gene and deletion of *exsA* did not affect phage plaque formation. (a) qRT-PCR analysis of *dap2* and *orf53* expression at 1-, 10-, 15-, and 30-min after phage infection. *dap2* was expressed immediately after entering the host and the expression decreased after 10 min. (b) The one-step growth curve of phage PaoP5 or PaoP5Δ*dap2* infecting PAO1 or Δ*exsA.* (a-b) Error bars indicate the mean ± SD (n=3) and statistical significance was determined using a two-sided Student’s *t-*test. (c) Phage PaoP5Δ*dap2* was mixed with PAO1 or Δ*exsA,* and used double-layer agar plates to observe the plaque number and size. Data are representative of three independent replicates.

**Fig. S5.**
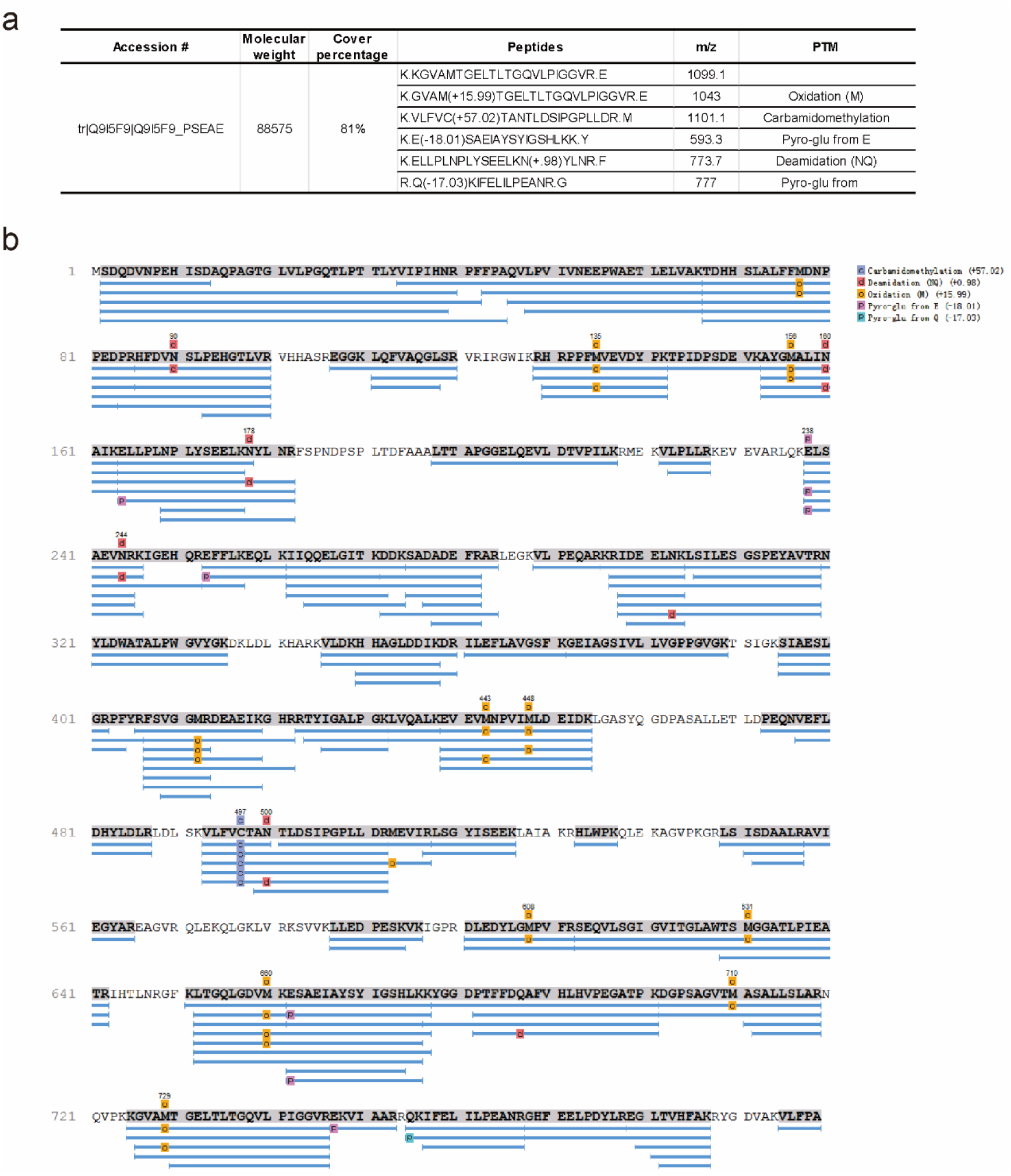
Identification of Lon protease that interacts with Dap2. (a) The specific protein bands identified in the Co-IP experiment were excised and subjected to mass spectrometry (MS) analysis. The MS results confirmed Lon as a novel interaction partner of Dap2, as indicated by the peptide matches and corresponding scores listed in the search results. (b) The coverage map displays peptide detection across the amino acid sequence of Lon, with peaks indicating identified regions.

**Fig. S6.**
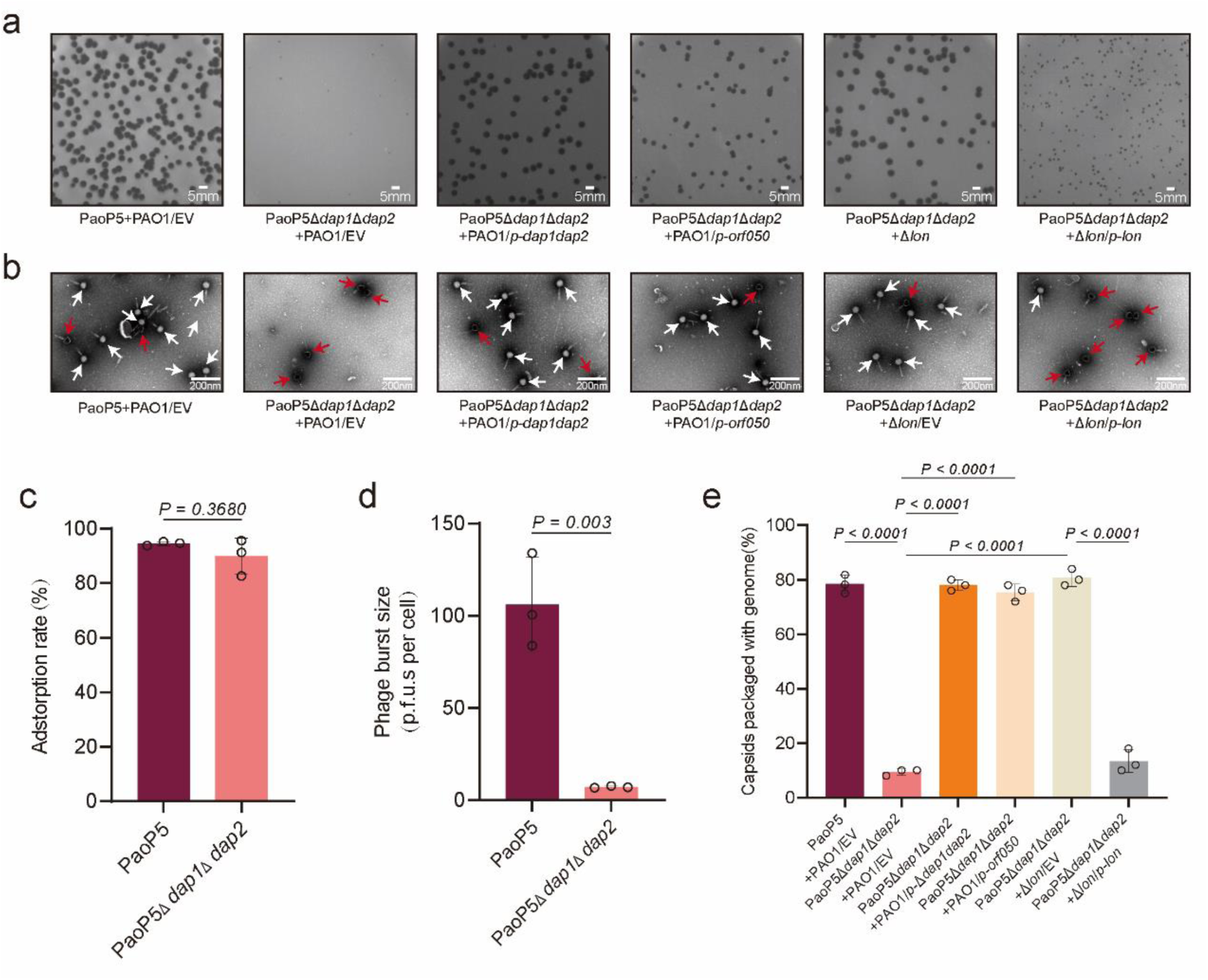
Impact of Lon and HNH on phage PaoP5Δ*dap1*Δ*dap2*. (a) The plaques of PaoP5 or PaoP5 *Δ dap1 Δ dap2* infecting PAO1/EV, PAO1/*p-dap1dap2,* PAO1*/p-hnh, Δlon*, and *Δlon/p-lon.* (d) Representative TEM of negatively stained phages produced in PAO1/EV, PAO1/*p-dap1dap2,* PAO1*/p-hnh, Δlon*, and *Δlon/p-lon.* The red arrows indicate the empty capsids and the phages in which the genome is packaged are indicated by the white arrows. (c) Both PaoP5Δ*dap1*Δ*dap2* and PaoP5 phages adsorbed to PAO1 efficiently. Error bars indicated the mean ± SD of four biological replicates. ns, no significance, based on Student’s *t* test. (d) The burst size of phage PaoP5 or PaoP5*Δdap1dap2* infecting PAO1. (e) Phages were cultured in the indicated strains, and the percentage of capsids packaged with genomes was calculated from three biological repeats. Three biological repeats were performed, and 50 particles were counted for the presence or absence of the genome. Error bars indicated the mean ± SD of three biological replicates. *P* value was calculated based on one-way ANOVA Dunnett’s multiple comparison test.

**Fig. S7.**
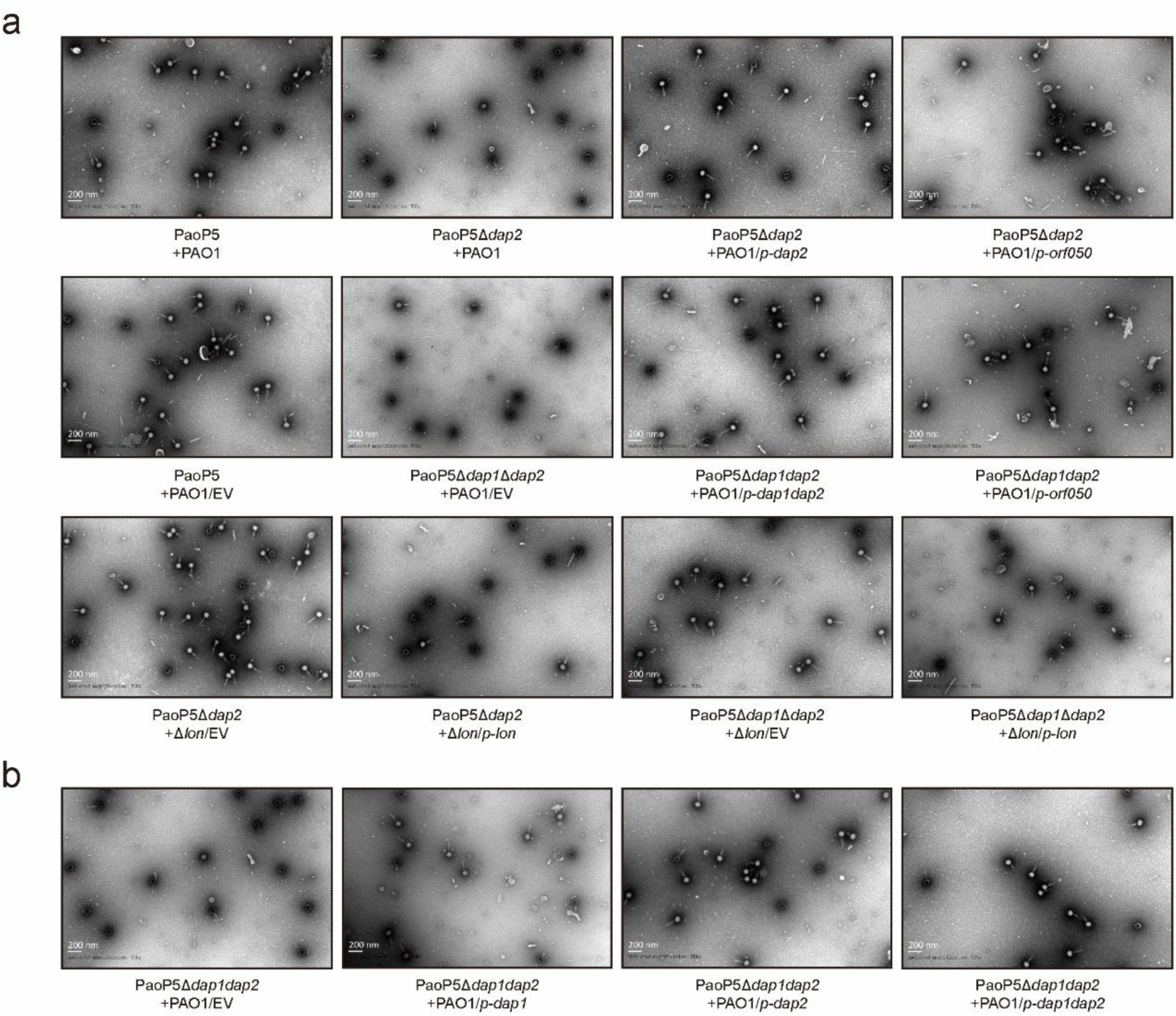
Representative transmission electron micrographs of negatively stained phages produced in PAO1, PAO1/*p-hnh*, PAO1/*p-dap2*, PAO1/*p-dap2dap2*, Δ*lon*, and Δ*lon*/*p-lon*. The empty capsids are black, and phages with white heads are packaged with the genomes.

## Materials and Methods

### Bacterial strains, phages, and culture conditions

The bacterial strains, phages, and plasmids used in this research are listed in Table S1. Both *E. coli* and *P. aeruginosa* strains were grown in Lysogeny Broth (LB) medium at 37°C with shaking at 220 rpm. Antibiotics were added as required at the following concentrations: for *E. coli*, carbenicillin (100 μg/mL), kanamycin (50 μg/mL), and tetracycline (10 μg/mL); for *P. aeruginosa* PAO1, carbenicillin (300 μg/mL), tetracycline (100 μg/mL), and gentamicin (50 μg/mL). Bacteriophages were propagated in LB medium at 37°C alongside their respective host bacteria.

### Construction of a promoter activity detection system

The promoterless *luxCDABE* reporter cluster in plasmid pMS402 was utilized to generate promoter-*luxCDABE* reporter fusions, following established methodology^28^. Target promoter regions were PCR-amplified and inserted into the pMS402 multiple cloning site. Recombinant plasmids were subsequently introduced into *Pseudomonas aeruginosa* PAO1 via electroporation. Bacterial luciferase activity, reflecting promoter-driven gene expression, was quantified in liquid cultures as light emission (counts per second, cps) using a Synergy 2 Multimode Microplate Reader (BioTek). Simultaneous measurements of luminescence and bacterial growth (optical density at 600 nm, OD_600_) were recorded at 30-minute intervals over a 24-hour incubation period. All data acquisition and analysis were performed using the instrument’s native software package.

### Construction of plasmids

To construct the pME6032-*dap2* plasmid, the DNA fragment was amplified by PCR using the primer pair *orf004-F* and *orf004-R* (Table S2). The PCR product was digested with EcoRI and HindIII restriction enzymes and then ligated into the similarly digested pME6032 vector, generating plasmid pME-*dap2*. This plasmid was transformed into the PAO1 strain, and transformants were selected on tetracycline-supplemented plates. Other plasmids were constructed using the same method.

To construct pHERD20T-*dap2* plasmid, the *orf004* gene was amplified using the primers pHERD-*orf004-F* and pHERD-*orf004-R*, and the resulting PCR product was ligated into the pHERD20T vector. The plasmid was then electroporated into the PAO1 strain, and transformants were selected on gentamicin-containing plates. The plasmid PAO1/p-*dap1dap2* was constructed similarly using the primers listed in Table S2. The plasmid pUCP24-dap2 was generated following the same procedure as described above.

### Western blot analysis

Overnight cultures of test strains were subcultured (1:100 dilution) in fresh LB medium supplemented with 5 mM EGTA and 20 mM MgCl_2_, then grown to mid-log phase (OD_600_ =0.6). Equal protein amounts (20 μg) were resolved by 12% SDS-PAGE and electrotransferred to PVDF membranes (Millipore). Membranes were blocked with 5% non-fat milk in TBST (50 mM Tris-HCl, 150 mM NaCl, 0.1% Tween-20, pH 7.6) for 1 h at 25°C, followed by overnight incubation at 4°C with mouse anti-Flag monoclonal antibody and rabbit anti-RNA polymerase α subunit as a loading control. After washing, membranes were incubated with HRP-conjugated goat anti-mouse IgG for 1 h at 25°C, and protein signals were visualized using an ECL Plus kit (Amersham Biosciences) on an ImageQuant LAS 4000 imager.

### GST Pull-down assay

In GST pull-down experiments, 100 μg of purified GST or GST-Dap2 was incubated with 25 μL pre-washed MagneGST™ glutathione beads (Promega) for 2 h at 4°C, followed by three washes with GST binding buffer to remove unbound proteins. Subsequently, 20 μg of His_6_-ExsA or His_6_-Lon was added to the bead complexes for 4 h at 4°C, after which nonspecific interactions were eliminated by washing with GST washing buffer. Retained proteins and input controls were resolved via SDS-PAGE, transferred to membranes, and subjected to immunoblotting using anti-GST (Immunoway) and anti-His (TransGen Biotech) antibodies, with protein signals visualized using an ECL Plus Kit (GE Healthcare, USA).

### RNA-seq and data analysis

RNA-seq was performed as previously described^36^. The PAO1/EV and PAO1/*p-dap2* strains were grown in LB medium at 37°C until the OD600 reached approximately 0.6. Total RNA was promptly isolated using TRIzol reagent (Invitrogen) following the manufacturer’s protocol. Ribosomal RNA was removed using the Ribo-Zero rRNA depletion kit. Subsequently, cDNA libraries were prepared and sequenced on an Illumina HiSeq 2500 platform. Each RNA-seq experiment was performed in triplicate. Sequencing reads were aligned to the *P. aeruginosa* reference genome (NC_002516.2, sourced from the National Center for Biotechnology Information) using Bowtie2, and only uniquely mapped reads were retained for further analysis. Differentially expressed genes (DEGs) were identified with DESeq2^37^, applying a Benjamini-Hochberg-adjusted P < 0.05 and |log2 fold change| > 1 as significance thresholds. The RNA-seq data have been deposited in the BioProject database under accession number PRJNA1232226.

### Real-time quantitative PCR (RT-qPCR)

To validate the RNA-seq data, PAO1 and PAO1/*p-dap2* samples were prepared as described above. To validate the expression of *dap2*, 10 mL of bacterial PAO1 culture (OD_600_=0.6) was infected with phage at an MOI of 10, and the culture grew at 37°C with shaking. For RNA extraction, 1 mL of the culture was taken at given time points. Three biological repeats were performed. Total RNA was extracted from each sample by RNAprep Pure Cell / Bacteria Kit (TIANGEN, China), rRNA was removed, and cDNA was generated using the PrimerScriptTM RT reagent kit with gDNA eraser (Takara, Japan). RT-qPCR was performed using 2X Universal SYBR Green Fast qPCR Mix (ABclone, China). The primers used in this study are listed in Table S1. The 16S rRNA gene was used as the reference gene for normalization, and the expression of each gene was compared using the delta-delta Ct method.

### Bacterial Two-Hybrid Assays

The bacterial two-hybrid assay was performed as previously reported^38^. Briefly, DNA fragments encoding Dap2 were PCR-amplified and cloned into the bait vector pBT to generate pBT-Dap2. The pTRG-ExsA vectors were constructed and co-transformed with pBT-Dap2 into the *E. coli* reporter strain. Transformants were selected on LB agar containing 5 mM 3-amino-1,2,4-triazole (3-AT) and incubated at 30°C for 48 h. Positive colonies were subsequently streaked onto dual-selection media supplemented with 5 mM 3-AT and 12.5 μg/mL streptomycin.

### Modelling for ExsA-Dap2 Complex Using Molecular Docking

The initial models of each protein with an average pLDDT values of ∼80 were generated by AlphaFold server^39^. Based on the predicted models, molecular docking was carried out to analyze the potential interaction modes between ExsA and Dap2, followed by one round of water refinement as implemented at the HADDOCK2.4 server [https://wenmr.science.uu.nl/haddock2.4/]^40,41^. The interaction interface of ExsA-Dap2 hetero-dimer candidate complex was further analyzed and evaluated by PDBePISA server [https://www.ebi.ac.uk/pdbe/pisa/]^42^. Carry incidentally, AF3 failed to predict the complex structure due to low ipTM value.

### Electrophoretic mobility shift assay (EMSA)

EMSAs were conducted as described^43^ to evaluate Dap2-ExsA interaction and its effect on ExsA DNA-binding activity. The *exsC* promoter region (amplified from *Pseudomonas aeruginosa* PAO1 genomic DNA) was incubated with an equimolar amount of His_6_-ExsA in binding buffer (20 mM Tris-HCl pH 7.5, 200 mM KCl, 2 mM EDTA, 2 mM DTT, 200 μg/mL BSA, 5 μg/mL salmon sperm DNA) for 10 min at 25°C, followed by addition of increasing concentrations of GST-Dap2 or GST-Dap2 point mutant for additional 10 min. Protein-DNA complexes were resolved on 6% native polyacrylamide gels in 0.5× TBE buffer (45 mM Tris-borate, 1 mM EDTA, pH 8.3) at 90 V for 1 h, stained with SYBR™ Gold nucleic acid gel stain (Invitrogen), and imaged using a Tanon-5500 phosphorimager. All assays were performed in triplicate with three biological replicates.

### Mouse infection and phage therapy experiment

The Animal Research Ethics Committee of Army Medical University reviewed, approved, and supervised all animal protocols (AMUMEC20250001). For the mouse infection study, PAO1/pUCP24 and PAO1/pUCP-*dap2* strains were cultured in LB medium at 37°C until reaching the early stationary phase. Bacterial cells were harvested, washed, and resuspended in PBS to an OD600 of 0.6. Each mouse group, consisting of 10 BALB/c female mice (7 weeks old), was intraperitoneally injected with 100 μL of bacterial suspension (∼2 × 10^7^ CFU) or PBS (negative control). Mice were monitored every 24 hours for7 days. Survivors at the end of the experiment were euthanized.

In the phage therapy experiment, 7-week-old BALB/c female mice were intraperitoneally injected with 50 μL of bacterial suspension (∼6 × 10^7^ CFU) or PBS (negative control). Two hours post-infection, 50 μL of phages (PaoP5, PaoP5*Δdap1*, or PaoP5*Δdap1Δdap2*, ∼6 × 10^8^ PFU) were administered intraperitoneally. Each group included 5 mice, which were monitored for 7 days. Survivors at the end of the observation period were euthanized.

To quantify phage and bacterial loads in the liver, measurements were taken up to 96 hours post-phage administration, as phages become undetectable after 4 days. A total of 32 mice were treated with PaoP5, and 80 mice were treated with PaoP5*Δdap1* or PaoP5*Δdap1Δdap2* to account for potential mortality during the experiment. Only surviving mice were used for quantification. At 1 h, 4 h, 8 h, 12 h, 24 h, 48 h, 72 h, and 96 h post-phage administration, 4 mice from each group were euthanized. Liver tissues were homogenized as previously described^22^. For phage titration, homogenized tissues were centrifuged at 10,000 × g for 10 minutes, and phage titers in the supernatant were determined using the EOP assay. Bacterial counts were obtained by spreading 100 μL of 10-fold serial dilutions of tissue samples onto LB agar plates in duplicate, followed by incubation at 37°C for 16 hours.

### Plaque assay and efficiency of plating (EOP) assay

The phage plaque assay was performed following established protocols^44^. Briefly, 200 μL of log-phase bacterial culture (OD600 of 0.6) was mixed with 100 μL of diluted phage solution (approximately 100 pfu) and 4 mL of 0.4% LB agar. The mixture was then overlaid onto LB agar plates and incubated overnight at 37 °C until plaques became visible.

To determine the efficiency of plating (EOP) of phages on different bacterial strains, a mixture of 4 mL of 0.4% LB agar and 200 μL of log-phase bacterial culture (OD600 of 0.6) was poured onto LB agar plates to create a double-layer agar. Subsequently, 2 μ L of serial 10-fold dilutions of the phage solution were spotted onto the prepared plates. The plates were incubated at 37°C for 18 hours to allow plaque formation.

### Knockout of phage genes using CRISPR-Cas9 system

The CRISPR-Cas9 plasmid pPTCS was utilized to knock out phage genes following established methods^43^. To delete *dap2* in phage PaoP5, a spacer was created by annealing the primers 004-G2-F/R (Table S2) and ligated into the Eco31I-digested pPTCS plasmid. For the recombination template, primers Δ004-LA-F/R and Δ004-RA-F/R were used to amplify two fragments upstream and downstream of the *dap4* gene, respectively. These fragments were ligated into the multiple cloning site of pTCPLS using Gibson Assembly. The resulting plasmid was introduced into PAO1 to generate PAO1/pTCPLS-Δg004G2D, which was then infected with 10^5^ PFU of phages. Survivors were selected using a plaque assay, and the mutant phage was confirmed by PCR and Sanger sequencing. The double deletion of *dap1* and *dap2* in phage PaoP5 was performed using the same approach, with the primers listed in Table S2.

### Bacteriophage adsorption assay

A phage adsorption assay was conducted as previously described^45^. Phages PaoP5*Δdap1Δdap2*, PaoP5*Δdap2*, or PaoP5 were mixed with *P. aeruginosa* PAO1 (OD600 of 0.6) at a multiplicity of infection (MOI) of 0.01. The mixture was incubated at 37°C for 10 minutes and then centrifuged. Phage titers in the initial phage solution (t1) and the supernatant (t2) were determined using the double-agar layer method. The adsorption rate was calculated using the formula (t1-t2)/t1. Statistical significance was assessed using Student’s t-test, with three biological replicates performed for each condition.

### One-step growth curve of phages

The one-step growth curve for PaoP5 *Δdap1Δdap2*, PaoP5 *Δdap2*, or PaoP5 was conducted as previously described^22^. Briefly, phages were mixed with 1 mL of log-phase bacterial culture at an MOI of 0.01 and incubated at 37°C for 10 minutes. The mixture was centrifuged at 10,000 × g for 1 minute, and the pellet was resuspended in 100 mL of LB medium. The suspension was incubated at 37°C for 60 minutes, with 0.2 mL of the culture sampled every 10 minutes. Each sample was centrifuged at 10,000 × g for 1 minute, and the phage titer in the supernatant was immediately measured using the EOP assay. The burst size was calculated as the ratio of the total number of phages released at the end of the growth cycle (40 minutes) to the total number of infected bacteria. Each experiment was performed in triplicate for every phage variant.

### Co-immunoprecipitation (Co-IP) assay

The GST Co-IP assay was performed as described previously^46^. Briefly, 100 μg of purified GST or an equal amount GST-Dap2 was mixed with 50 μL of pre-washed MagneGST glutathione beads slurry (Progma) in GST binding buffer (50 mM Tris, 150 mM NaCl, pH 8.0) and incubated for 2 h at 4°C, after which unbound GST proteins were washed away. Then, the PAO1 cell lysates were incubated with the protein-binding beads for an additional 6 h at 4°C. The beads were then washed with GST washing buffer (50 mM Tris, 500 mM NaCl, pH 8.0) sufficiently and the beads bound proteins were dissolved in SDS sample buffer. After SDS-PAGE, proteins were visualized by silver staining (Beyotime, China). The samples from the gels (molecular weight from 26 to 100 kDa.) retained by beads coated with GST-Dap2, but not GST alone, were analyzed by Matrix-assisted laser desorption/ionization (MALDI) mass spectrometry.

### Transmission Electron Microscopy (TEM)

The phages PaoP5*Δdap1Δdap2*, PaoP5*Δdap2*, or PaoP5 were propagated on various bacterial strains, including PAO1, PAO1/*p-orf050*, PAO1/*p-dap2*, PAO1/*p-dap1dap2*, *Δlon,* or *Δlon/p-lon*. The phage lysates were analyzed using transmission electron microscopy (TEM) following a standard protocol^22^. Briefly, the lysate was applied to carbon-coated copper grids for 10 minutes, followed by negative staining with a 2% phosphotungstic acid solution for 30 seconds. Phage particles were then examined using TEM, and the percentage of phages with genome-containing capsids was determined by averaging counts from over 50 particles across three independent biological replicates.

### Protein expression and purification

To construct the pGEX-6p-1-*dap2, pGEX-6p-1-dap2^V52K^,* pGEX-6p-1-*dap2*^A64K^, pGEX-6p-1-*dap2*^N79A^, pGEX-6p-1-*dap2*^V52KA64KN79A^, pGEX-6p-1-*dap2*^E97AE99A^, *pET28a-dap1*, *pET28a-hnh*, and *pET28a-lon* plasmids, the fragments of *dap1, dap2, hnh,* and *lon* were amplified by PCR with the corresponding primers listed in Table S2, respectively. Subsequently, the amplified fragment was digested with corresponding enzymes and ligated into pET28a, or pGEX-6p-1 vector, respectively. The resultant plasmids were introduced into *E. coli* BL21(DE3). After this, the strains were cultured at 37°C in LB medium. When the OD_600_ reached 0.6, 0.5 mM isopropyl β-D-1-thiogalactopyranoside (IPTG) was added to induce protein production. The cultures were incubated at 16°C for an additional 20 hours.

The cells were collected via centrifugation and resuspended in buffer A [20 mM tris-HCl (pH 8.0), 300 mM NaCl, and 25 mM imidazole]. The cells were lysed by high-pressure homogenization and centrifuged. The supernatant was applied to a 5-mL HisTrap HP column (Cytiva), and the target protein was eluted via AKTA purifier (GE Healthcare) using buffer B [20 mM tris-HCl (pH 8.0), 300 mM NaCl, and 500 mM imidazole]. The target protein was collected and applied to a HiLoad 16/600 Superdex 75-pg gel filtration column (Cytiva) equilibrated with buffer composed of 20 mM tris-HCl (pH 8.0), 200 mM NaCl, and 2 mM dithiothreitol (DTT). The purified proteins were concentrated and stored at −80°C.

### Protein degradation assay

The HNH endonuclease degradation assay was carried out as previously described^47^. Briefly, 100 μM of Dap2 and/or 100 μM of Dap1 were combined with 100 μM of HNH endonuclease and incubated on ice for 20 minutes. The reaction mixture was then added to 30 μg of Lon in a 50 μL buffer containing 4 mM ATP, 50 mM Tris-HCl (pH 8.0), 10 mM MgCl_2_, 1 mM DTT, 80 μg/mL creatine phosphokinase (Sigma, #C3755), and 50 mM creatine phosphate (Sigma, #27920). The mixture was incubated at 37°C for the specified duration, and a 10 μL aliquot was collected at each time point. SDS-PAGE loading buffer was added to the aliquot, followed by heating at 100°C for 15 minutes. The degradation of HNH was analyzed using 12% SDS-PAGE. The extent of HNH protein degradation was assessed as described above.

### Phylogenetic analysis

Homologs of *dap1* and *dap2* were identified by searching against 26,648 phage genomes retrieved from the INPHARED database on January 20, 2025. The search was performed using MMseqs2^48^, and hits with an e-value below 10^-5^ were retained. Sequence alignment and phylogenetic trees were conducted using MEGA7^49^, and the resulting trees were visualized using iTOL^50^.

### Statistical analyses

The statistical analysis employed in this study involved the utilization of the student’s *t*-test to compare data from two distinct groups. A significance level of *P < 0.05* was adopted to determine statistical significance.

### Data Availability

The analyzed data and raw RNA-seq readings of PAO1 expressing *dap2* were uploaded to the NCBI GEO (PRJNA1232226).

