## Supplemental Table 1 for "Bacteriophage protein Dap2 inhibits bacterial type III secretion system and synergizes with Dap1 to evade anti-phage immunity"

Table S1. Strains plasmids and phages used in the study

| **Strain or plasmid** | **Relevant characteristics** | **Source** |
| --- | --- | --- |
| ***P. aeruginosa*** |  |  |
| PAO1 | Reference strain, derivative of PAO | (Liang et al., 2008) |
| Δ*exsA* | *exsA* knockout mutant of PAO1; Gm^r^ | (Kong et al., 2013) |
| Δ*lon* | *lon* deletion mutant of PAO1 | (Yang et al., 2015) |
| Δ*lon*/*p-lon* | *lon* complement strain，  derived from Δ*lon* and pAK1900-*lon*;Cb^r^ | (Yang et al., 2015) |
| **Phage** |  |  |
| PaoP5 | *P.aeruginosa* phage | (Shen et al., 2016) |
| PaoP5Δ*dap2* | Knockout *orf004* in PaoP5 | This study |
| PaoP5Δ*dap1*Δ*dap2* | Knockout *orf014* in PaoP5 | This study |
| PaoP5Δ*orf014* | Knockout *orf014* in PaoP5 | This lab |
| PAO1/pME6032-*dap2* | Overexpression of *orf004* using pME6032 plasmid in PAO1 | This study |
| PAO1/pUCP24-*dap2* | Overexpression of *orf004* using pUCP24 plasmid in PAO1 | This study |
| PAO1/*p*-*dap1* | Overexpression of *orf003* using pHERD20T plasmid in PAO1 | This lab |
| PAO1/*p*-*dap2* | Overexpression of *orf004* using pHERD20T plasmid in PAO1 | This study |
| PAO1/*p*-*dap1dap2* | Overexpression of *orf003orf004* using pHERD20T plasmid in PAO1 | This study |
| PAO1/*p-orf050* | Overexpression of *orf050* using pHERD20T plasmid in PAO1 | This lab |
| ***E. coli*** |  |  |
| DH5α | *F – φ80lacZ ΔM15 Δ(lacZYA-argF)U169 recA1 endA1 hsdR17(rk– , mk+)phoA supE44 thi-1 gyrA96 relA1 tonA* | Stratagene |
| BL21（DE3） | *F-, ompT, hsdSB (rBB - mB - ), gal, dcm ( DE3 )* | Stratagene |
| BTH101 | *F-, cya-99, araD139, galE15, galK16, rpsL1 ( Strr ), hsdＲ2, mcrA1, mcrB1* | (McCarthy et al., 2017) |
| XL1-BlueMRF' Kan H | Host strain for propagating pBT and pTRG recombinants;*Δ(mcrA)183Δ(mcrCB-hsdSMR-mrr)173 endA1 supE44 thi-1 recA1 gyrA96 relA1 lac* [F' *proAB lacI^q^ Z ΔM15* Tn*5*(Kn^r^)] | Agilent |
| XL1-BlueMRF' Kan R | Report strain of BacterioMatch II two-hybrid system;*Δ(mcrA)183Δ(mcrCB-hsdSMR-mrr)173 endA1 supE44 thi-1 recA1 gyrA96 relA1 lac* [F' *laqI^q^ HIS3 aadA* (Kn^r^)] | Agilent |
| **Phage** |  |  |
| PaoP5 | *P.aeruginosa* phage | (Shen et al., 2016) |
| PaoP5Δ*dap2* | Knockout *orf004* in PaoP5 | This study |
| PaoP5Δ*dap1*Δ*dap2* | Knockout *orf003orf004* in PaoP5 | This study |
| PaoP5Δ*orf014* | Knockout *orf014* in PaoP5 | (Liang et al., 2024) |
| **Plasmids** |  |  |
| pME6032 | Plasmid for over-expression of gene in PAO1 and PaoP5 | (Heeb et al., 2002) |
| pMMB67EH-Flag | pMMB67EH vector with Flag tag coding sequence | (Chen et al., 2020) |
| pHERD20T(Gm^R^) | Plasmid for over-expression of gene in PaoP5 | (Zhou et al., 2022) |
| pUCP24 | Plasmid for over-expression of gene in PaoP5 | (Lasarte-Monterrubio et al., 2022) |
| pUT18C | Plasmid for bacterial-two hybrid assays | (McCarthy et al., 2017) |
| pKT25 | Plasmid for bacterial-two hybrid assays | (McCarthy et al., 2017) |
| pET28a | Plasmid with His tag for protein expression and purification | (Wang et al., 2020) |
| pTCPLS | Plasmid with CRISPPR-cas9 for gene knockout in PaoP5 | (Yang et al., 2023) |
| pMS402 | Plasmid with lux-based reporters for Luminescence screening assays | (Duan et al., 2003) |
| pBT | p15A origin of replication, *lac-UV5* promoter, λ cI open reading fram; Cm^r^ | Agilent |
| pTRG | ColE1 origin of replication, lpp promoter, lac-UV5 promoter, RNAPα open reading frame; Tc^r^ | Agilent |
| pBT-LGF2 | Interaction control plasmid encoding the dimerization domain (40 amino acid residues) of the Gal4 transcriptional activator protein; Cm^r^ | Agilent |
| pTRG-GAII 1^p^ | Interaction control plasmid encoding a domain (90 amino acid residues) of the mutant form of the GaII 1 protein; Tc^r^ | Agilent |
| pKD-*exsA* | pMS402 containing *exsA* promoter region; Kn^r^, Tmp^r^ | (Duan *et al.*, 2003) |
| pKD-*exsC* | pMS402 containing *exsCEB* promoter region; Kn^r^, Tmp^r^ | (Duan *et al.*, 2003) |
| pKD-*exoS* | pMS402 containing *exoS* promoter region; Kn^r^, Tmp^r^ | (Duan *et al.*, 2003) |
| pKD-*exoT* | pMS402 containing *exoT* promoter region; Kn^r^, Tmp^r^ | (Duan *et al.*, 2003) |
| pKD-*exoY* | pMS402 containing *exoY* promoter region; Kn^r^, Tmp^r^ | (Duan *et al.*, 2003) |
| mini-CTX*-exsA*(*p_exsC_*)-FLAG | Mini-CTX-*lacZ* containing the entire *exsA* gene driven by *exsCEBA* promoter fused with FLAG tag at C-terminal; Tc^r^ | This lab |
| pGEX-6p-1 | Expression vector with N-terminal GST tag; Amp^r^ | This lab |
| pGEX-6p-1-*dap2* | pGEX6p-1 derivative for expression of *dap2* | This study |
| pGEX-6p-1-*dap2*^V52K^ | pGEX6p-1 derivative for expression of *dap2*^V52K^ | This study |
| pGEX-6p-1-*dap2*^A64K^ | pGEX6p-1 derivative for expression of *dap2* ^A64K^ | This study |
| pGEX-6p-1-*dap2*^N79A^ | pGEX6p-1 derivative for expression of *dap2* ^N79A^ | This study |
| pGEX-6p-1-*dap2*^V52KA64K N79A^ | pGEX6p-1 derivative for expression of *dap2* ^V52KA64K N79A^ | This study |
| pGEX-6p-1-*dap2*^E97AE99A^ | pGEX6p-1 derivative for expression of *dap2* ^E97AE99A^ | This study |
| pET28a-*exsA* | Protein expression construct, the entire gene of *exsA* cloned in pET28a vector | This study |
| pET28a-*dap1* | Protein expression construct, the entire gene of *dap1* cloned in pET28a vector | (Liang et al., 2024) |
| pET28a-*hnh* | Protein expression construct, the entire gene of *orf050* cloned in pET28a vector | This study |
| pET28a-*lon* | Protein expression construct, the entire gene of *lon* cloned in pET28a vector | This study |
| pBT-*dap2* | pBT plasmid containing the entire *dap2* gene | This study |
| pTRG-*exsA* | pTRG plasmid containing the entire *exsA* gene | This study |
| pUT18C-*dap2* | *dap2* cloned into pUT18C for complementation | This study |
| pKT25-*dap1* | *dap1* cloned into pKT25 for complementation | This study |
| pKT25-*hnh* | *hnh* cloned into pKT25 for complementation | This study |
| pKT25-*lon* | *lon* cloned into pKT25 for complementation | This study |
