## Supplemental Table 2 for "Bacteriophage protein Dap2 inhibits bacterial type III secretion system and synergizes with Dap1 to evade anti-phage immunity"

**Table S2: Primers used in this study**

| **Primers** | **Sequence (5’–3’)** |
| --- | --- |
| **Overexpression of**  **genes in PAO1** |  |
| *p*-*orf004*-F | GCGAATTCGAGCTCGGTACatgtacgacaaggctcaag |
| *p*-*orf004*-R | AGCTTGCATGCCTGCAttagttttcattctcttcctccattt |
| *p*-*orf003004*-F | AAACGATGGCGATTGCGatgtacgacaaggctcaagt |
| *p*-*orf003004*-R | CGACGGCCAGTGCCAttagaccctcccgaac |
| pME6032-*orf004*-F | CAAgaattcatgtacgacaaggctcaa |
| pME6032-*orf004*-R | CCGctcgagttagttttcattctcttc |
| pUCP24- *orf004*-F | ACGAATTCGAGCTCGGTACatgtacgacaaggctca |
| pUCP24- *orf004*-R | TGCCTGCAGGTCGACTttagttttcattctcttcctcc |
| **Knock out phage genes** |  |
| *004-*G1F | TAGTacacttcatgagtgtgacac |
| *004*-G1R | AAACgtgtcacactcatgaagtgt |
| Δ*004*-LA-F | GGTCTGACAGCTCGAGagtctaatgagaactgcttcc |
| Δ*004*-LA-R | agttttcatgaagccgtttccgc |
| Δ*004*-RA-F | aacggcttcatgaaaactaaagagatcgatgtc |
| Δ*004*-RA-R | TTTTTTTGGCGCGCCattgctcagacattagaccc |
| Δ*003004*-LA-F | GGTCTGACAGCTCGAGagtctaatgagaactgcttc |
| Δ*003004*-LA-R | ctcccgagaagccgtttccgctc |
| Δ*003004*-RA-F | acggcttctcgggagggtctaatgtc |
| Δ*003004*-RA-R | TTTTTTTGGCGCGCctccctagtggcgtagttc |
| Δ*003*-RO | gcgatggttgtatagattcacc |
| Δ003-LO  Δ004-RO | agacccctgaagagaacatc  gcttttgatgcacccgatag |
| Δ*004*-LO | tccttacgcccgaaaaag |
| **3-AT** |  |
| pBT-004-F | ATAgcggccgcAATGTACGACAAGGCCTCAA |
| pBT-004-R | CACgaattcTTAGTTTTCATTCTCTTCCTCCATT |
| pTRG-*exsA*-F | ATAgcggccgcAATGCAAGGAGCCAAATAT |
| pTRG-*exsA*-R | CACgaattcTCAGTTATTTTTAGCCCGGC |
| **BATCH** |  |
| pUT18C-*004*-F | TGGAACGCCACTGCAatgtacgacaaggctcaagt |
| pUT18C-*004*-R | AGTGCACCATATTACTTAGTTATATttagttttcattctcttcctccattt |
| pKT25-*003*-F | ACGCGGCGGGCTGCAatgaaaactaaagagatcgatgt |
| pKT25-*003*-R | AGTGAATTCTTACTTACTTAGGTACttagaccctcccgaacgtt |
| pKT25-*050*-F | ACGCGGCGGGCTGCAatgaagctgtgccctcgc |
| pKT25-*050*-R | AGTGAATTCTTACTTACTTAGGTACctaaactacaatctcatatccttcaatacc |
| pKT25-*lon*-F | ACGCGGCGGGCTGCAatgaaaacactcgtcgaattg |
| pKT25-*lon*-R | AGTGAATTCTTACTTACTTAGGTACctaatgcgtgctaattcgc |
| **RT-qRCR** |  |
| 16s-F | CAAAAGCTACTGAGCTAGAGTACG |
| 16s-R | TAAGATCTCAAGGATCCCAACGGCT |
| *exsA*-qpcr-F | GGAGAATCCTCTATGCCCATCA |
| *exsA*-qpcr-R | CTCTGGGTGAAATAGGACTGACTG |
| *exsC*-qpcr-F | TGGCACCGTTTCGATCTGCA |
| *exsC*-qpcr-R | GCCAAGGTCGCCTCGAAGCATT |
| *exoS*-qpcr-F | AGGAGCTGGATGCGGGACAAA |
| *exoS*-qpcr-R | CCACGGGTGCCACGGAAAGT |
| *exoT*-qpcr-F | CGCGAAATCGCCGTCCAA |
| *exoT*-qpcr-R | AGCCCGAAGTGCTCCACCAG |
| *exoY*-qpcr-F | GCATGGCAGTGGTGGTCTCG |
| *exoY*-qpcr-R | CCATAGAATCCGTCCTCGCTCA |
| *hemH*-qpcr-F | TGGAAAGCGTGCGTCCGTACCTG |
| *hemH*-qpcr-R | GGGCTGACGTTGCGGCTGTTCT |
| *hfp*-qpcr-F | CACGCAAGCAGGCGGAGAGC |
| *hfp*-qpcr-R | TGGCAGCGGAAAGCGAATAG |
| *pscF*-qpcr-F | GAATACCCTCGATACCGTGG |
| *pscF*-qpcr-R | GTTGATGTTGTAGATGACCG |
| *pcrV*-qpcr-F | CCGAAGCAGAGCGGGGAA |
| *pcrV* -qpcr-R | CCGAGTTGTAGCGGGAGC |
| *orf004*-qpcr-F | AGTGCTACTGTGGGTTGG |
| *orf004*-qpcr-R | TGGCCTCTTTGAGTTCTT |
| *orf053*-qpcr-F | CGGTTCTGTCGGTGGTCT |
| *orf053*-qpcr-R | TCAACAGGCTCGTCGTCT |
| **Protein expression and purification** |  |
| pEGX-6p-1-*orf004*-F | CTGTTCCAGGGGCCCatgtacgacaaggctcaa |
| pEGX-6p-1-*orf004*-R | CGTCAGTCAGTCACGATGCttagttttcattctcttcctccatttct |
| *orf004-*DTB-F | cgggtgtactaccgagactta |
| *orf004*-DTB-R | cttgcttggctagggttgct |
| *orf004*^V52K^-F | acctacgacttccgaaggaAAgacttccatttcgtca |
| *orf004*^V52K^-R | tgacgaaatggaagtcTTtccttcggaagtcgtaggt |
| *orf004*^A64K^-F | gagaagaaagtgaccgAAtacacacttcatgagtgtga |
| *orf004*^A64K^-R | tcacactcatgaagtgtgtaTTcggtcactttcttctc |
| *orf004*^N79A^-F | gtcctactacagagctaGCtggcacctacacgcttca |
| *orf004*^N79A^-R | tgaagcgtgtaggtgccaGCtagctctgtagtaggac |
| *orf004*^E97AE99A^-F | ggaagCgaatGCAaactaaagagatcgatgtc |
| *orf004* ^E97AE99A^-R | gttTGCattcGcttcctccatttctgcaa |
| **Promoter** |  |
| *excC* promoter-F | GCCGTCTCCGCGCGGGAGGA |
| *excC* promoter-R | GGGGGCGCCTCCTAAAGCTC |
